# maskNMF: A denoise-sparsen-detect approach for extracting neural signals from dense imaging data

**DOI:** 10.1101/2023.09.14.557777

**Authors:** Amol Pasarkar, Ian Kinsella, Pengcheng Zhou, Melissa Wu, Daisong Pan, Jiang Lan Fan, Zhen Wang, Lamiae Abdeladim, Darcy S. Peterka, Hillel Adesnik, Na Ji, Liam Paninski

## Abstract

A number of calcium imaging methods have been developed to monitor the activity of large populations of neurons. One particularly promising approach, Bessel imaging, captures neural activity from a volume by projecting within the imaged volume onto a single imaging plane, therefore effectively mixing signals and increasing the number of neurons imaged per pixel. These signals must then be computationally demixed to recover the desired neural activity. Unfortunately, currently-available demixing methods can perform poorly in the regime of high imaging density (i.e., many neurons per pixel). In this work we introduce a new pipeline (maskNMF) for demixing dense calcium imaging data.

The main idea is to first denoise and temporally sparsen the observed video; this enhances signal strength and reduces spatial overlap significantly. Next we detect neurons in the sparsened video using a neural network trained on a library of neural shapes. These shapes are derived from segmented electron microscopy images input into a Bessel imaging model; therefore no manual selection of “good” neural shapes from the functional data is required here. After cells are detected, we use a constrained non-negative matrix factorization approach to demix the activity, using the detected cells’ shapes to initialize the factorization. We test the resulting pipeline on both simulated and real datasets and find that it is able to achieve accurate demixing on denser data than was previously feasible, therefore enabling faithful imaging of larger neural populations. The method also provides good results on more “standard” two-photon imaging data. Finally, because much of the pipeline operates on a significantly compressed version of the raw data and is highly parallelizable, the algorithm is fast, processing large datasets faster than real time.

## Introduction and overview

Calcium and voltage imaging enable in vivo recording from large populations of neurons. A major goal of these methods is to simultaneously record the activity of as many neurons as possible at high spatial and temporal resolution. A number of approaches have been developed in pursuit of this goal; see e.g. (Ji et al., 2016) for a review.

One promising strategy is to optically mix the activity of many cells onto the image sensor — to effectively increase *R*_*c*_, the ratio of cells per pixel — then computationally demix the activity after the images have been acquired. Examples include compressed sensing approaches (Pnevmatikakis and Paninski, 2013), Bessel imaging (Lu et al., 2017), the vTwINS approach developed in Song et al. (2017), the two-stage imaging approach proposed in Friedrich et al. (2017a), and the multiplexing approach in (Yang et al., 2016). Ideally, we want to make *R*_*c*_ as large as possible to increase the size of the neural population that we can image simultaneously (under the constraint that demixing remains feasible). However, the denser the imaging, the more challenging the resulting computational demixing problem — and unfortunately, current demixing approaches can fail on dense neural data in which there is a high spatial overlap.

To address this issue, we have designed a pipeline that sequentially compresses and temporally sparsens calcium imaging video data. Then we deploy a specialized neural network to detect neural shapes from the resulting video. This neural network is trained on simulated calcium imaging data constrained by electron microscopy, to sidestep the need to gather large labeled datasets from human observers that may inject uncontrolled biases into the training dataset.

## Background and related work

### Segmentation and matrix factorization approaches

Over the past decade, a variety of calcium and voltage imaging signal extraction methods have been developed. Broadly speaking, we can divide these analysis approaches into two classes: *segmentation* methods and *matrix factorization* methods. Segmentation methods aim to define clear, non-overlapping region of interest (ROI) boundaries for each neuron and assign the activity in each ROI pixel to at most a single neuron (Pachitariu et al., 2013; Kaifosh et al., 2014; Apthorpe et al., 2016; Klibisz et al., 2017; Spaen et al., 2019; Kirschbaum et al., 2020; Bao et al., 2021); to extract activity of any neuron from the video then we simply need to perform a weighted average over the pixels belonging to its ROI. On the other hand, matrix factorization methods are more general, and allow for contributions from multiple cells to each pixel (Mukamel et al., 2009; Ferran and Hamprecht, 2014; Maruyama et al., 2014; Pnevmatikakis et al., 2016; Pachitariu et al., 2016; Petersen et al., 2018; Zhou et al., 2018; Buchanan et al., 2018; Zhou et al., 2020; Inan et al., 2017; Reynolds et al., 2017; Charles et al., 2022). The goal of a matrix factorization method is therefore to *demix* the activity of multiple cells that may have some partial spatial overlap. These demixing methods are more appropriate than segmentation methods for the highly-mixed data we consider in this paper.

Within this class of matrix factorization approaches, constrained or penalized nonnegative matrix factorization (NMF) methods have been widely applied (Ferran and Hamprecht, 2014; Maruyama et al., 2014; Pnevmatikakis et al., 2016; Pachitariu et al., 2016; Inan et al., 2017; Zhou et al., 2018; Buchanan et al., 2018; Zhou et al., 2020). Here the data matrix is factorized into a sum of non-negative product terms: each neuron’s activity is modeled as a product of a fixed non-negative spatial image multiplied by a time-varying non-negative intensity factor. This is a natural generative model of the observed imaging data (after preprocessing steps such as motion correction / registration have been applied), but unfortunately computing the optimal NMF solution can be a challenging computational problem.

NMF involves approximating a solution to a nonconvex optimization problem. Therefore, most NMF algorithms are highly initialization-dependent. There are known mathematical conditions that ensure a good solution to the NMF problem: for example, if it is known that, for each neuron *i*, there exists a pixel that only contains signal from neuron *i* (the “pure-pixel” assumption), or conversely, there exists a frame that only contains signal from neuron *i* (the “pure-frame” assumption), then existing algorithms can detect either pure pixels or pure frames and use these to initialize an NMF solution that correctly identifies the visible neurons. (See (Arora et al., 2016) for an example of such mathematical guarantees.) Unfortunately, in the dense imaging case that we focus on here, these pure-pixel or pure-frame assumptions do not hold.

The key insight of this paper is that if we can sparsen the data while maintaining a low noise level, then a local version of the pure-frame assumption is reasonable: well-isolated neurons are visible in many frames in the sparsened, denoised data. In the calcium imaging setting, we can sparsen the data with a temporal deconvolution operation (together with a suitable denoising operation), and then use a specialized neural network to detect the resulting isolated cell images. (See (Xie et al., 2020) for a related approach, in which high-pass thresholding sparsens voltage imaging videos.) Finally, we use the detected cell images to initialize a constrained NMF algorithm to demix the full original dense data.

### Neural network based approaches

Recent years have seen the development of multiple calcium imaging pipelines that use deep neural networks (Apthorpe et al., 2016; Klibisz et al., 2017; Soltanian-Zadeh et al., 2019; Kirschbaum et al., 2020; Zhou et al., 2020; Cai et al., 2021; Denis et al., 2020). In some approaches, networks are trained to extract solely spatial information. For example, in (Klibisz et al., 2017), a neural network is trained to detect neural shapes from a video’s mean image. This approach exposes the network to a 2D “summary” image, essentially discarding most temporal information. However, this approach has the advantage of being simple and computationally fast. In other approaches, networks are trained to use spatiotemporal information (Apthorpe et al., 2016; Soltanian-Zadeh et al., 2019). For example, (Soltanian-Zadeh et al., 2019) uses a specialized 3D convolutional neural network to extract neural signals. The network makes more complete use of the spatiotemporal structure of calcium imaging videos. However, it is also computationally more expensive. One hybrid approach uses both spatial and temporal statistics to train a convolutional neural network. This network estimates affinities between pixels of a given imaging video (Kirschbaum et al., 2020). These affinities are then processed to estimate individual neural signals. Training these neural networks poses a challenge, as there is no ground truth in calcium or voltage imaging videos. Instead, previous efforts have mainly relied on manually annotated datasets (Apthorpe et al., 2016; Klibisz et al., 2017; Soltanian-Zadeh et al., 2019; Kirschbaum et al., 2020). Unfortunately, these annotations are not universally accepted, even among experts (Giovannucci et al., 2019; Soltanian-Zadeh et al., 2019).

We take a different approach here: we use simulated calcium imaging videos as training data, constraining our simulation models with real morphological information extracted from electron microscopy datasets (Zhou et al., 2020), as discussed in more depth below. (See (Charles et al., 2019) for a different, fully-artificial approach to simulating calcium imaging data.)

### Denoising and compression approaches

Another class of methods use deep neural networks for calcium imaging analysis tasks such as noise reduction or neuropil estimation (Lecoq et al., 2021; Zhou et al., 2020; Zhang et al., 2022). (Lecoq et al., 2021) builds on a line of work (Lehtinen et al., 2018; Batson and Royer, 2019; Krull et al., 2018) to create a self-supervised deep neural network denoiser for functional neuroimaging data. Additionally, Zhang et al. (2022) extends the simulation approach from Charles et al. (2019) to train a deep neural network to remove neuropil contamination from one photon imaging data, making it easier to detect ROIs. These image-to-image denoising approaches are complementary to the approach taken in this paper; for example, one could apply one of these denoising networks first and then apply the demixing pipeline described below.

In this work, we adapt the Penalized Matrix Decomposition method from (Buchanan et al., 2018) to compress and denoise the data. This step estimates a *signal subspace* directly from the data (without having to pre-train a neural network) and discards noise dimensions orthogonal to this signal subspace, therefore denoising the data.

In addition, the data can be reprojected onto this signal subspace iteratively at multiple stages of the processing pipeline, helping to e.g. suppress noise after the temporal sparsening step across pixels. Finally, by projecting onto the signal subspace we achieve significant compression, leading to major speedups of subsequent algorithm steps, as we will describe below.

## Methods

### Notation and overview

We will use the following notation, following (Buchanan et al., 2018):

1. *Y* refers to the full “unrolled” imaging video, formed by vectorizing each frame and stacking these vectors into a matrix. *Y* is a *d* × *T* matrix, where *T* is the number of frames in the video and *d* is the number of pixels in the field of view (FOV).
2. *N* refers to the number of neurons identified in the video *Y* .
3. *A* denotes the the spatial footprints of the *N* neurons. It is a *d* × *N* matrix.
4. *C* denotes the temporal activities of the *N* neurons. It is a *N* × *T* matrix.
5. *B* describes a “background” term; this is a *d* × *T* matrix.
6. *E* is a noise term, a *d* × *T* matrix. We assume that noise is temporally and spatially uncorrelated.
7. 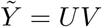 refers to the rank-*K* compressed and denoised representation of *Y*, where *U* is a *d* × *K* matrix and *V* is a *K* × *T* matrix (discussed below).

Putting these pieces together, we use the following model:

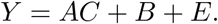

Our goal is to generate estimates of *A, C, B*, and *E*. Broadly, our pipeline can be broken into three stages: preprocessing, initialization and demixing. During the preprocessing stage, we aim to register the data and compress it. The former step spatially aligns the signal sources across the frames of the movie. The latter step reduces the data size (allowing for faster subsequent data processing) and also serves as a conservative denoiser. During the initialization stage, we aim to create good initial estimates of *A* and *C*. This step is particularly challenging in dense imaging datasets, where neurons have a high degree of overlap. Finally, during the demixing stage, we use an improved version of the “localNMF” demixing procedure developed in Buchanan et al. (2018) to estimate *A, C*, and *B*.

Below we describe the proposed pipeline in greater detail. Note that every step in this pipeline is designed for GPU acceleration, allowing for end-to-end analysis results for hour-long imaging videos within minutes.

### Motion correction and registration

We begin by motion correcting the video using NoRMCorre (Pnevmatikakis and Giovannucci, 2017).

Conceptually, NoRMCorre is a template matching algorithm: it aligns every frame of the video to a “global” template. This alignment involves a “rigid” registration, wherein the entire image is uniformly shifted to best match the template, as well as an optional “piecewise rigid” step, wherein local patches of the image are aligned to the corresponding subpatches of the template. These aligned patches are then pieced together via interpolation to arrive at a final registered image. For computational performance and parallelization, we have implemented the entire algorithm on GPU using JAX (Frostig et al., 2018). This allows us to rapidly perform multiple iterations of rigid and piecewise rigid registration for any dataset to estimate an excellent template to which we register the data.

### Compression and Denoising

After motion correction, we next compress and denoise the registered, normalized video to produce a new video 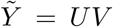 following (Buchanan et al., 2018). In this decomposition, *U* represents a compressed spatial representation of the signal, and *V* represents a compressed temporal representation. We use a simple version of the penalized matrix decomposition (PMD) method described in (Buchanan et al., 2018), without the spatial and temporal penalties, to compute *U* and *V*. This is equivalent to applying adaptive, localized singular value decompositions (SVDs) to overlapping spatial subsets of the movie (and then linearly interpolating over these spatial subsets to avoid block artifacts). The rank *K* of *U* and *V* is typically an order of magnitude or two smaller than *d* or *T* ; additionally, each column of *U* is supported only on a single spatial patch, and therefore *U* is a highly sparse matrix. This step is highly parallelized over spatial video patches and is again accelerated for GPU and multi-GPU environments using JAX (Frostig et al., 2018). Furthermore, using JAX’s composability mechanisms, we provide an option to perform end-to-end motion correction and compression on the GPU without ever explicitly saving out a motion corrected movie. This approach is further accelerated by JAX’s just-in-time compilation mechanism, and can be very useful in time-constrained (online) experimental settings. Finally, we note that unlike the original PMD method, we use a fast approximate method from (Halko et al., 2011) to compute the truncated SVD; this method relies largely on matrix-vector operations, enabling fast GPU acceleration.

### Sparsening

Highly-overlapping populations of neurons are difficult to demix. In the calcium imaging setting, the observed fluorescence signals are temporally much slower than the underlying action potentials: this spreads the activity over multiple imaging frames, leading to greater spatially-mixed data in each frame.

Our approach is to temporally deconvolve the data, producing a sparser dataset. Deconvolving the raw data in each pixel would be challenging, due to the low SNR in each individual pixel. Instead, we deconvolve the temporal activity of each pixel in the denoised video 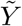, by solving the deconvolution problem described in OASIS (Friedrich et al., 2017b):

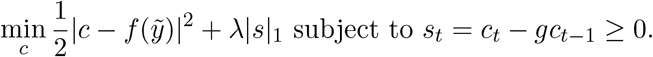

In the context of deconvolution, 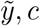 and *s* are *T* -length vectors (one for each pixel) describing the fluorescence trace, estimated calcium concentration, and deconvolved calcium trace, respectively. The function *f* subtracts the baseline fluorescence from 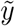 and *s* are related via an AR-1 process with coefficient *g*. The regularization coefficient *λ* encourages sparsity in the underlying activity *s*. (It is worth noting that other deconvolution methods could also work well here, e.g., (Jewell and Witten, 2018; Berens et al., 2018; Jewell et al., 2019; Rupprecht et al., 2021).)

The above approach requires *O*(*dT*) time, and the current implementation of OASIS does not take advantage of GPU architectures. To achieve further speedups, we can apply a more naive weighted deconvolution for an AR-1 process:

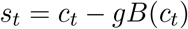

where B is the backshift operator applied to the time series *c*. This simpler deconvolution operation is less accurate when applied to noisy data but has a major advantage: it can operate exclusively on the temporal basis representation (*V*) of the denoised data (leading to *O*(*KT*) scaling rather than *O*(*dT*)) and is highly parallelizable via standard GPU matrix manipulation. This yields significantly more scalable computation, especially over large-FOV data.

### Re-denoising via U-Projection

In the simplest case, deconvolution is a differentiation operation, and therefore amplifies high temporal-frequency noise in the data. The OASIS model employed above reduces this noise amplification, but since this deconvolution is run on each pixel independently (for code simplicity and speed) some noise persists in each frame. Therefore, next we perform a linear projection of the deconvolved matrix *Z* back onto the spatial denoising matrix *U*, to recover a cleaner representation of the deconvolved activity:

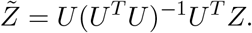

To compute this projection efficiently, we first calculate the inverse matrix (*U*^*T*^ *U*)^−1^. This operation is relatively quick because *U*^*T*^ *U* is a low-dimensional, *K* × *K* matrix, with *K << d*. Then we simply need to multiply by *U*^*T*^ and *U* ; both of these are quick due because *K* is small and *U* is sparse (Buchanan et al., 2018). The projected video 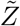 typically contains isolated, clean neuron shapes, which we can detect using a Mask R-CNN object detection network (see the next section). Finally, we note that for purposes of memory efficiency, we never explicitly compute 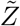and store it in memory; instead we leave it in a factorized form: 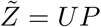 where *P* = (*U*^*T*^ *U*)^−1^*U*^*T*^ *Z*. In Figs. 2-3 we provide pixel-wise and frame-wise examples to show how the deconvolution and projection operations improve the isolation and detection of neural signals.

**Figure 1:**
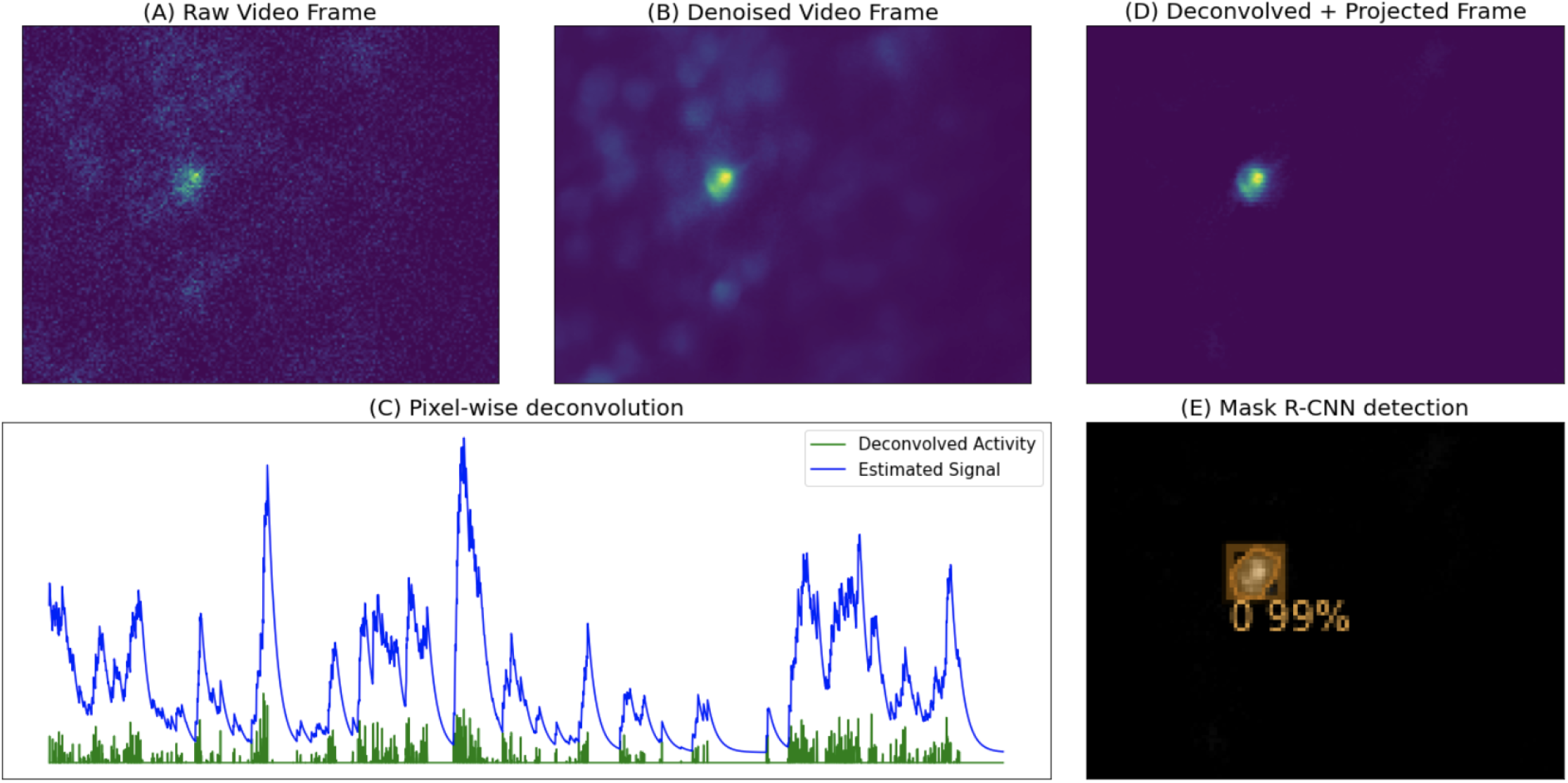
Illustrated steps of maskNMF pipeline. (A) We start with a motion-corrected dataset. (B) We then compress and denoise the data using penalized matrix decomposition (PMD) from Buchanan et al. (2018). (C) Then, we perform pixel-wise deconvolution to temporally sparsen the data; here, we show the result of temporally sparsening a single pixel of the denoised data. (D) We then project this sparse video back onto the spatial denoising basis *U* ; note that the resulting image contains a single, well-isolated neural shape, unlike (B). (E) Finally, we run a specialized neural network (a Mask R-CNN architecture, trained on simulated calcium imaging data) to detect neuron signals present in this frame of the data.

**Figure 2:**
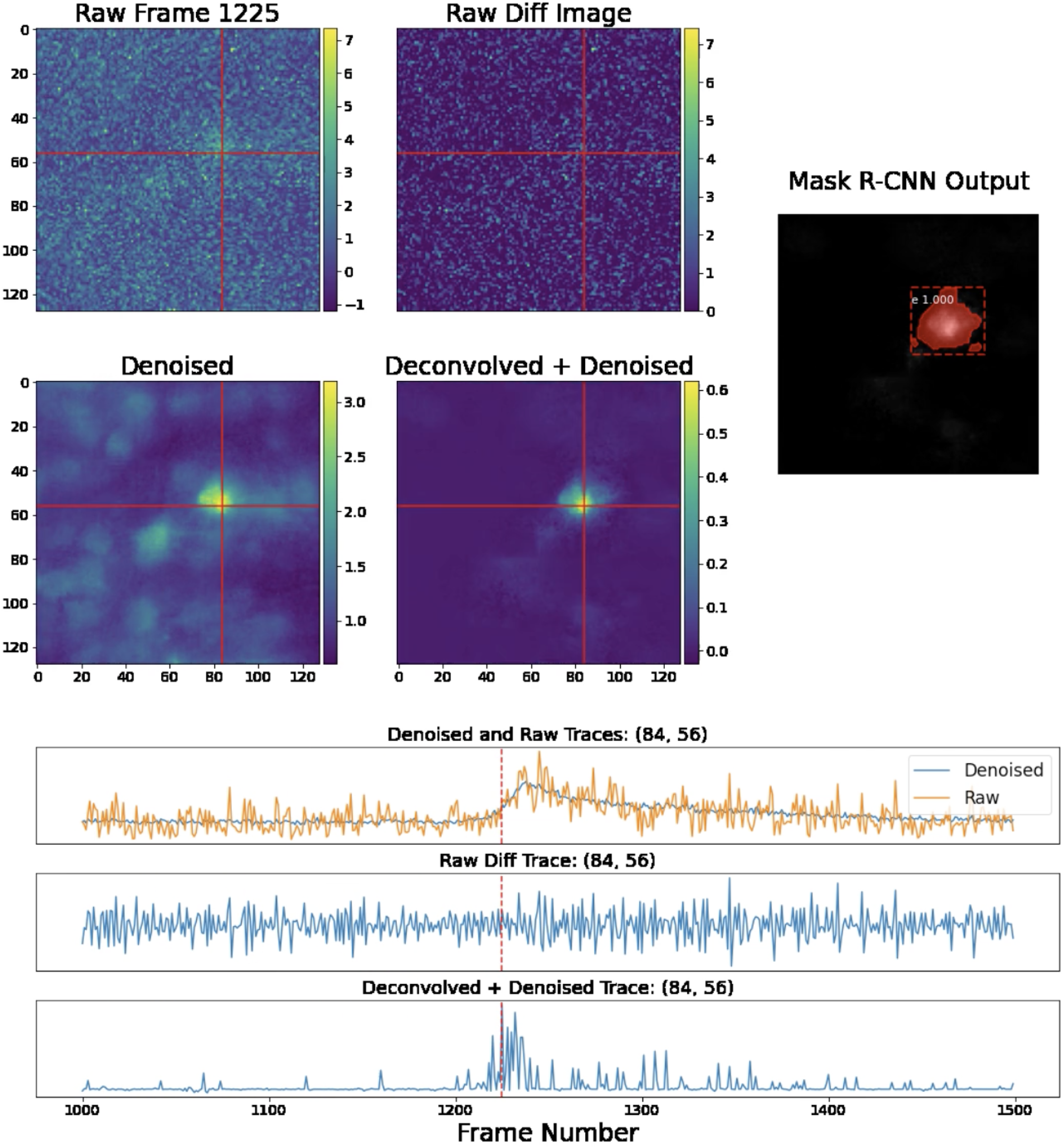
Single-frame and single-pixel comparisons on Bessel data. We provide comparisons of various images and the temporal traces in a fixed pixel, (84,56), as indicated by the crossed lines. The original and denoised image highlight the effects of the first denoising step in this pipeline. Following this step, we deconvolve and denoise the temporal traces of each pixel, producing the projection frame. We provide the difference image as a simple baseline with which to compare the projection; the difference operator acts as a very crude deconvolution here. The difference image simply calculates the difference between frame 1225 and 1226; note that this raw difference is highly noise-contaminated. Finally, the projection frame is much sparser than the denoised frame, making it easier to detect spatial footprints of individual neurons, which we do using Mask R-CNN. For further details, see the full corresponding video [here](https://drive.google.com/file/d/1wN_GTE0krNRXlCwjcSS1ngIRDUJJG1eZ/view?usp=sharing).

**Figure 3:**
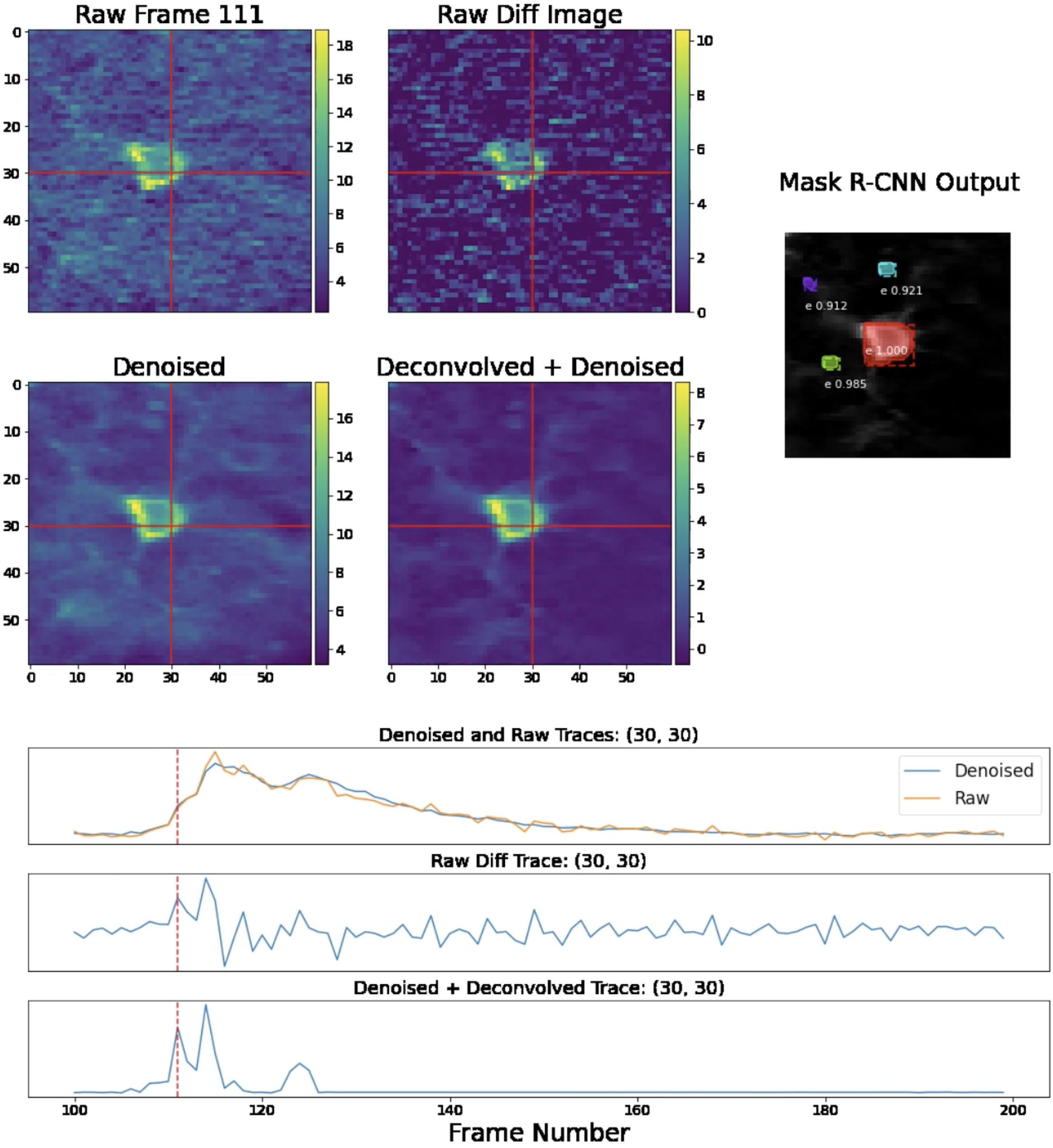
Single-frame and single-pixel comparisons for standard two-photon data. We provide comparisons of various images and the temporal traces for two-photon data in the same fashion at Fig. 2. Note that the Mask R-CNN network shown here is specialized to detect neuron shapes found in non-volumetric two-photon data. Again, see the full video here for further details.

**Figure 4:**
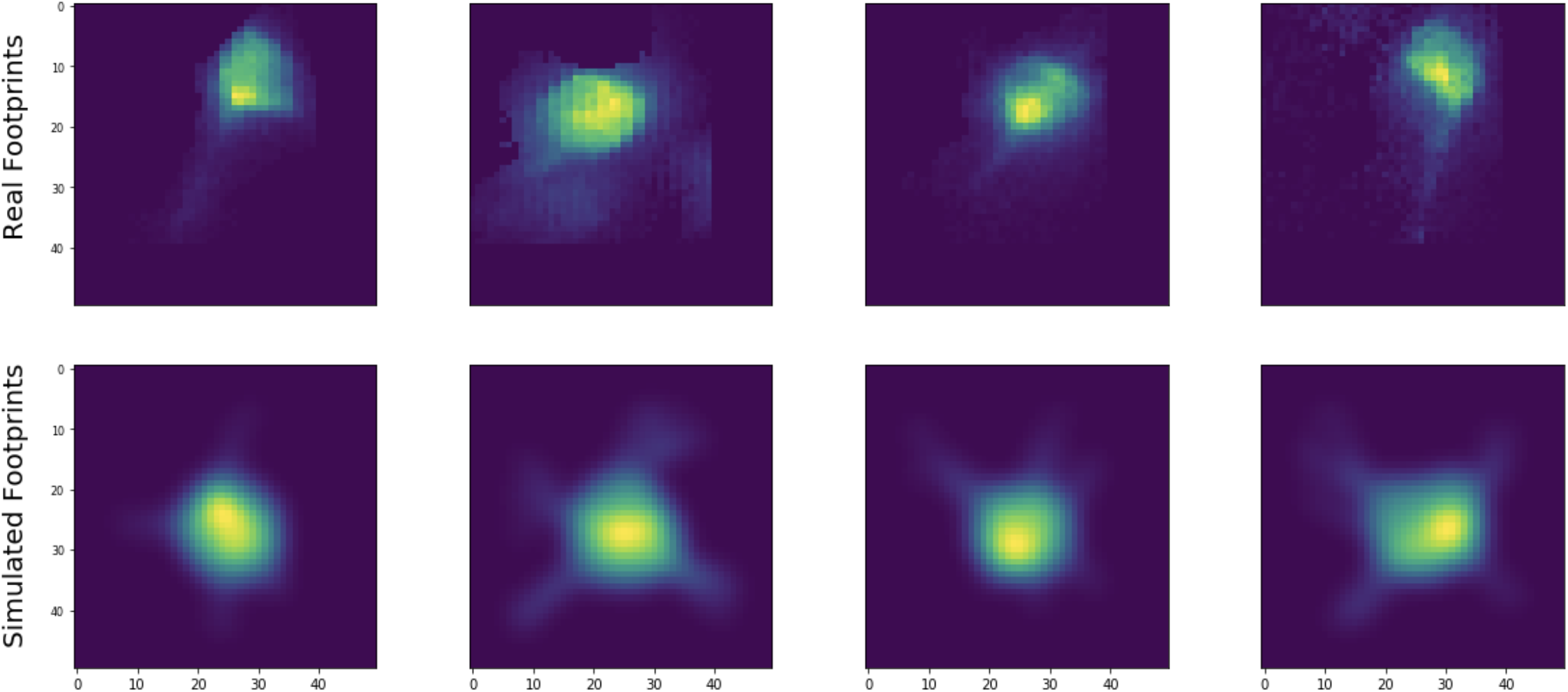
Simulated versus real spatial footprints. Examples of neural shapes extracted from real data using the proposed pipeline (top) versus simulated neural shapes used to train the detection network (bottom). The simulated neuron somas and processes are reasonably well-calibrated to those of the real neurons.

**Figure 5:**
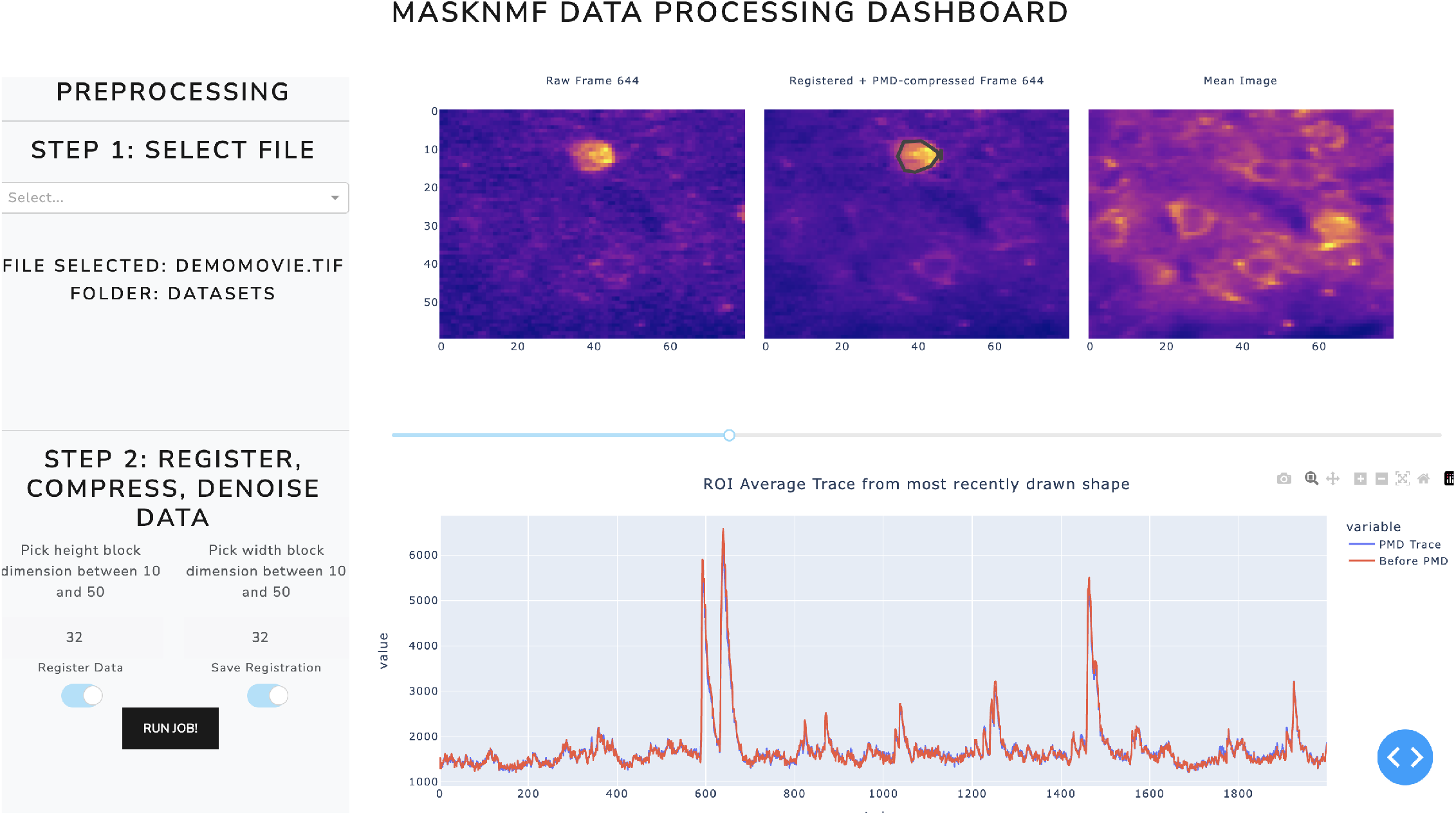
Dash web-app user interface. Screenshot of the motion correction and PMD compression stages of the user interface.

### Component identification: Mask R-CNN

Now, we use an object segmentation network, Mask R-CNN (He et al., 2017), to identify candidate neurons by their spatial supports in 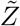 This network is trained in a supervised manner (see the next section for details) to perform an *instance* segmentation task. That is, it learns to identify individual instances of a certain class (in this case, neurons) given images that may contain neurons. (We use Mask R-CNN because it is easy to train using our simulated neuron shapes; however, other methods, such as (Dolev et al., 2019), might also work well here.) The brightest frames in 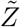 are more likely to contain the desired complete neuron images. To construct a reliable initialization for *A*, we reorder the frames of the movie 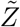 based on maximum brightness in each frame. We run Mask R-CNN on the brightest *n* frames (with *n* ≈ 100), creating a list of candidate neurons. The Mask R-CNN identifies neurons with varying levels of confidence. We only add a candidate neuron to our list if it meets certain criteria. First, the Mask R-CNN must identify the neuron with a confidence level of more than *c*_*min*_ (with *c*_*min*_ ≈ 0.7). Second, the cosine similarity between the candidate neuron and all other neurons already in the list must be lower than some threshold, *t*_*real*_. Third, the cosine similarity between the candidate neuron’s spatial support and the spatial supports of all neurons already in the list must be lower than some threshold, *t*_*bin*_. We have found *t*_*bin*_ ∈ [0.5, 0.8] and *t*_*real*_ ∈ [0.5, 0.8] work well on a variety of datasets. These criteria allow us to construct a list of high-confidence, distinct neurons to initialize *A*. Finally, the candidate neuron footprint must not overlap significantly with any other footprints in its respective frame. This last condition allows us to identify well-isolated, non-overlapping neural signals. To seed *A*, we provide the spatial footprints of the neurons in the candidate list.

See Table 1 for a summary of the hyperparameters used in this section and the next.

**Table 1:** Mask R-CNN detection and training hyperparameters.

| Variable | Description | Recommended Range |
| --- | --- | --- |
| $c_{min}$ | Confidence threshold for Mask R-CNN | [0.7] |
| $t_{real}$ | Footprint cosine similarity threshold | [0, 5, 0.8] |
| $t_{bin}$ | Binary mask cosine similarity threshold | [0.5, 0.8] |
| $n$ | # of frames of $\tilde{Z}$ on which we run Mask R-CNN for each local region | [50, 150] |
| $m$ | Max # of neuron footprints in simulated video | dataset dependent |
| $p$ | Percentile threshold for deconvolved traces | [40, 80] |

We run the above initialization procedure in parallel on overlapping local spatial patches. (For reference, we have found that using a roughly 20 × 20 pixel patch is appropriate for all of the datasets analyzed here, where a typical soma occupied between 100-300 pixels.) This has several advantages. First, if the brightness varies significantly across a given video’s FOV, then simply taking the brightest components across the entire image would lead us to neglect cells from the dimmer parts of the field of view. Second, recall that we iteratively build a candidate list of neurons, disregarding neurons that are similar to neurons already in the list. The runtime of this operation is quadratic in the number of candidate cells; running this operation locally keeps the size of this list (and therefore the runtime of this step per patch) constant as the size of the full FOV increases.

### Video simulation and training

We train the Mask R-CNN network using simulated data. Specifically, we generate simulated *A, C*, and *E* matrices, form a simulated data matrix *AC* + *E* and compute the corresponding 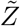 and finally compute the *n* brightest frames in local spatial patches, following the steps outlined above. This simulated training strategy enables us to generate an unlimited amount of training data without any laborious manual labeling of individual cells in real video data. On the other hand, the accuracy of the resulting trained network will depend heavily on the realism of the simulated training data. We note that the proposed training procedure is highly modular: if desired, we can use a different simulation method to generate the neuron spatial profiles included in the *A* matrix. To generate accurate neural shapes in the simulated *A* matrices, we simulate Bessel imaging spatial footprints using electron microscopy (EM) data. (Zhou et al., 2020) describe a method for simulating two-photon calcium imaging data from a library of three-dimensional segmented EM neural shapes. In this work, we expand on that method and use a dataset-dependent point-spread function to simulate training data. For example, for Bessel imaging data we simulate neural shapes using a Bessel point-spread function instead of a standard two-photon (Gaussian) scanning point spread function; this has the effect of collapsing the three-dimensional EM shape over the length of the Bessel beam in *z*.

To simulate a 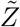 video, we choose a random number *k* and then place *k* neurons, each multiplied by a random scaling factor (to account for variations in brightness between cells), onto a common field of view. (The distribution of the number of cells *k* is also dataset-dependent: to match denser datasets, we can sample a larger number of cells per field of view.) For each neuron *i*, we randomly select a temporal trace *c*_*i*_ to populate the temporal activity matrix *C*. To train a Mask R-CNN for Bessel data analysis, we used “real” Bessel traces from other source extractions; however, we expect that simulated calcium traces would also work well. We then construct the raw video *Y* = *AC* + *E*, where *E* represents Gaussian white noise; following Buchanan et al. (2018), we let the variance of *E* scale with the size of the signal *AC* in each pixel.

For each frame in the resulting simulated video, we must decide which neurons should be considered as “positive” (i.e., which neurons have an activity level that the Mask R-CNN should detect after sufficient training). To do so, we individually deconvolve the temporal traces in *C* and soft threshold, keeping only temporal activity in the top *p*-th percentile. (We found values in the range of *p* ∈ [40, 80] work well and allow for quick training convergence.) This defines a new matrix of thresholded temporal traces *C*^′^. In a given frame, the Mask R-CNN is expected to identify neuron *i* if its corresponding thresholded temporal trace in *C*^′^ is nonzero at that frame.

We used the implementation of Mask R-CNN provided by Wu et al. (2019). We used default parameters, including standard learning rates (0.2). We generated approximately 400 different simulated videos to use as training data.

### Demixing denoised data: localNMF

As discussed above, we use the Mask R-CNN to identify a set of candidate neurons, and we initialize *A* = *A*_*init*_ with the neuron shapes provided by Mask R-CNN. To initialize the temporal activity terms *C* = *C*_*init*_, we perform a hierarchical alternating least squares (HALS) (Cichocki et al., 2007) update, performing non-negative regression of *A*_*init*_ onto the denoised video, 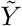 Then to further improve our estimates of *A, C* and *B* we run an updated version of the localNMF algorithm from (Buchanan et al., 2018), using an improved version of the ring background model adapted from (Zhou et al., 2020) (discussed further below). In the appendix, we provide detailed explanations of our enhancements to the original localNMF procedure; in particular, the new approach performs all computations on GPU and operates on the *U, V* decomposition rather than the original large data matrix *Y*, leading to major acceleration.

### Simplified background model

We model the background *B* as the sum of a fluctuating, time-varying background term and a static background term, given by *B*_*f*_ and *B*_*s*_ respectively.

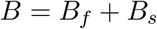

At each stage of the localNMF demixing procedure, we define the static background term, *B*_*s*_, to be the mean of the residual *UV* − *AC*.

To model *B*_*f*_, we adapt the ring background model from CNMF-E (Zhou et al., 2018), with two modifications. First, any ring pixel belonging to the support of *A* is set to zero. Second, the ring weights for any pixel are constant on their support. This drastically reduces the number of free parameters in the ring model, leading to faster fitting and reduced overfitting while still achieving accurate background subtraction. Finally, to denoise *B*_*f*_, we perform a linear subspace projection of *B*_*f*_ onto *U* in the same manner as we describe in the “U-projection” section above. Again, performing these computations on GPU, and in the *U, V* subspace rather than on the full data matrix *Y*, both lead to major accelerations.

### Superpixels and multi-pass analysis

In the localNMF demixing procedure, we include a multi-pass approach to identify and demix any neural signals which may be left in the residual *UV* − *AC*. To do so, we run a more robust version of the superpixels initialization procedure from (Buchanan et al., 2018) to identify the remaining neural signal; see the appendix for a detailed explanation.

### Dockerized Dash app and Napari viewer

We provide a Dash application via a Docker container which allows users to rapidly and interactively motion correct, compress, and demix large calcium imaging data. This app contains a detailed set of visualization options which will allow experimentalists to quickly inspect their data, annotate it, generate high quality demixing results, and download the results in a convenient format for downstream analysis tasks; see documentation [here](https://masknmf-full-pipeline.readthedocs.io/en/latest/), including instructions for cloud execution.

Dash apps are not designed for video rendering. We know, however, that inspection of final processed images/movies is a critical component of data inspection, so we have created an export feature and a Napari plugin to efficiently view these compressed movies on a local workstation (Napari, 2021).

## Results

### Timing

We have run this pipeline on a variety of real and simulated calcium imaging datasets. We provide approximate timing measurements on an imaging dataset of dimensions 512 × 512 × 50, 000 (about 30 minutes at 30 Hz), on a 128GB RAM Ubuntu workstation with a NVIDIA GeFORCE GPU with 11GB RAM. These numbers are useful to give a coarse sense of the relative timing of each of the steps described above.

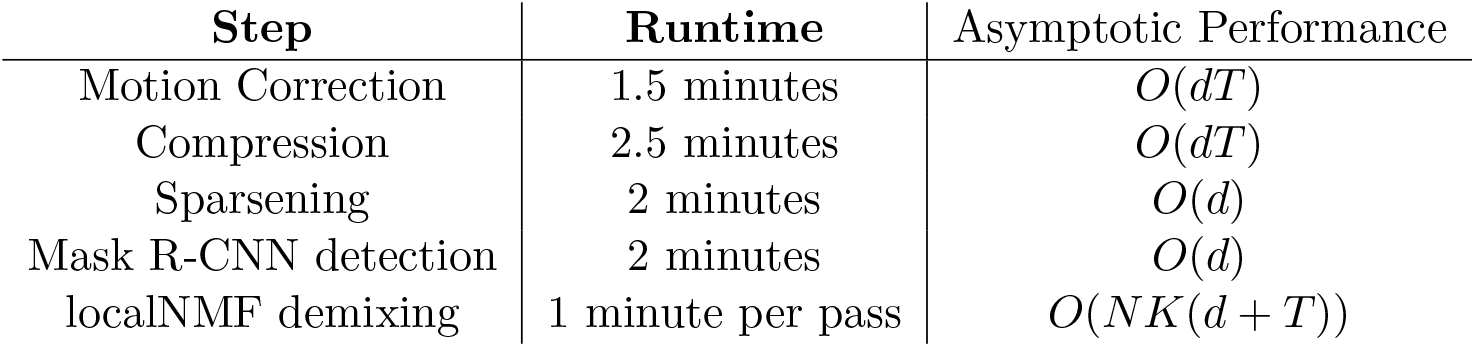

Thus, the entire pipeline takes minutes and operates significantly faster than real time. We note that the localNMF demixing only takes one minute. This is made possible by the PMD step: we can fit the entire compressed data, (*U, V*) onto the GPU and run the entire localNMF demixing pipeline there. Since most operations in this demixing step involve matrix multiplications, this leads to massive acceleration.

### Simulation evaluation

We first evaluate the performance of the new pipeline on simulated data, where ground truth is available. We generate more than 200 simulated videos across a wide range of neuron densities and score the performance of the new pipeline along with two baseline algorithms (discussed below). We follow the video simulation procedure outlined above: we simulate a set of spatial footprints, randomly place them onto a common FOV, sample a corresponding set of temporal traces, and generate *Y* = *AC* + *E*, a noisy background-free simulated Bessel imaging movie.

Fig. 6 provides a sample frame of a simulated *demixing video*, a useful diagnostic that gives a side-by-side view of all key aspects of our pipeline, from the raw data, to the denoised outputs, background estimates, and residual.

**Figure 6:**
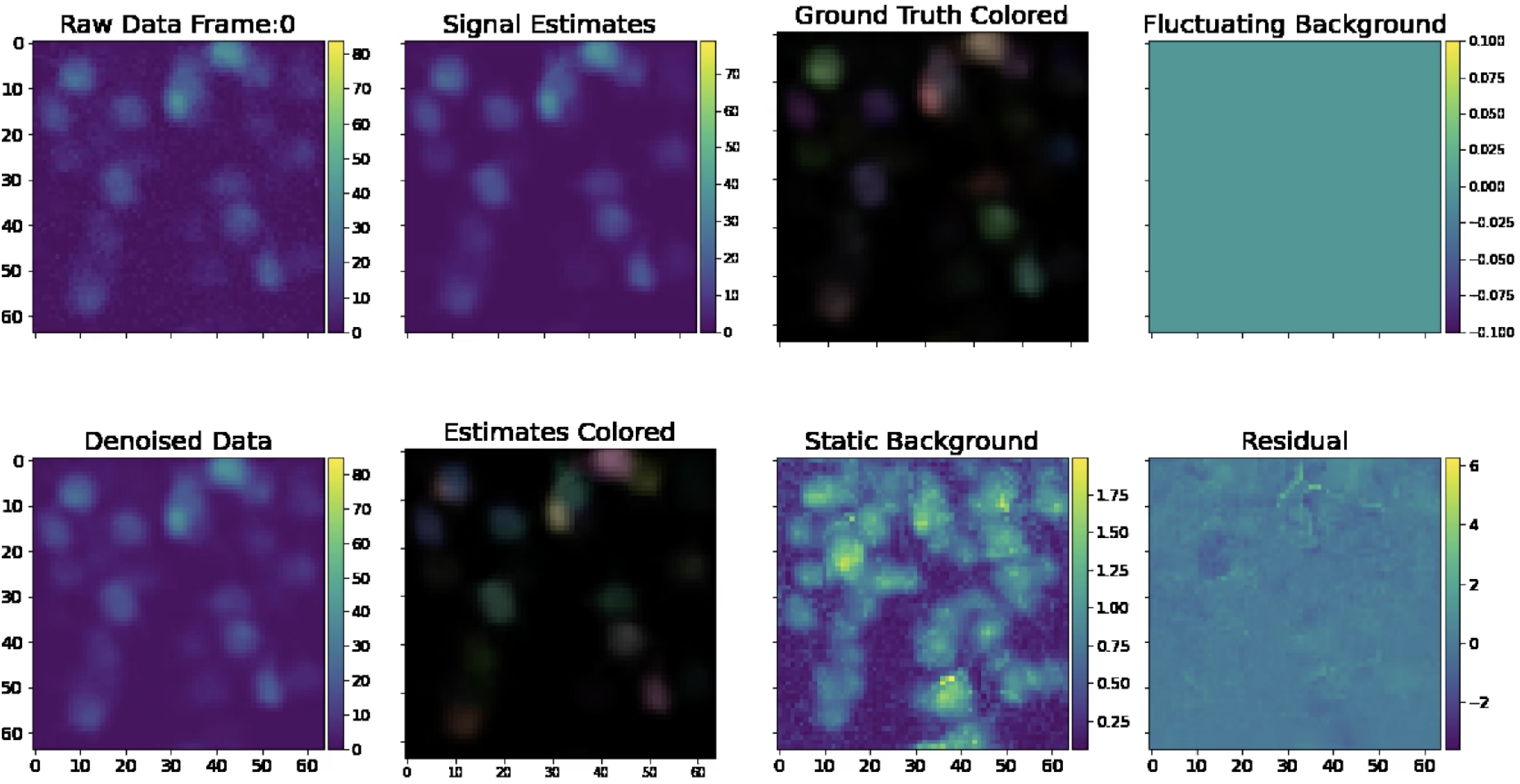
Sample frame of a simulated demixing video. Here, we provide a single frame in a demixing video, which displays all key aspects of our pipeline. The “Raw Data” frame displays the data after noise normalization (so the estimated noise level in each pixel is one). The “Denoised Data” frame describes the PMD-denoised data. The “Signal Estimates” describe the summed estimated spatial and temporal components *AC*. In the “Estimates Colored” panel, we assign each estimated component *a*_*i*_*c*_*i*_ an individual color so that the viewer can visually discern nearby neurons. In the “Ground Truth” frame, we provide the ground truth neural activity in the same fashion. We provide static background estimates in the “Static Background” panel. We provide a “Fluctuating Background” panel for reference. Note that it is empty because our simulated data does not contain neuropil activity. Finally, we provide a mean-subtracted residual frame in the “Mean Sub Residual” panel. The residual is calculated by subtracting the signal and neuropil estimates from the PMD-denoised movie. The full demixing video is available here.

The simulated dataset shown here has density *R*_*c*_ = 2.4. Visual comparison of the ground truth and recovered components here indicates high recovery accuracy; in addition, the residual frame, which is calculated by subtracting the signals and neuropil from the denoised movie, indicates that most of the signal in the video has been captured accurately. Finally, in Fig. 7, we provide a library of all components extracted from an example spatial subpatch of the field of view of Fig. 6. The new pipeline accurately identifies the spatial and temporal components present here.

**Figure 7:**
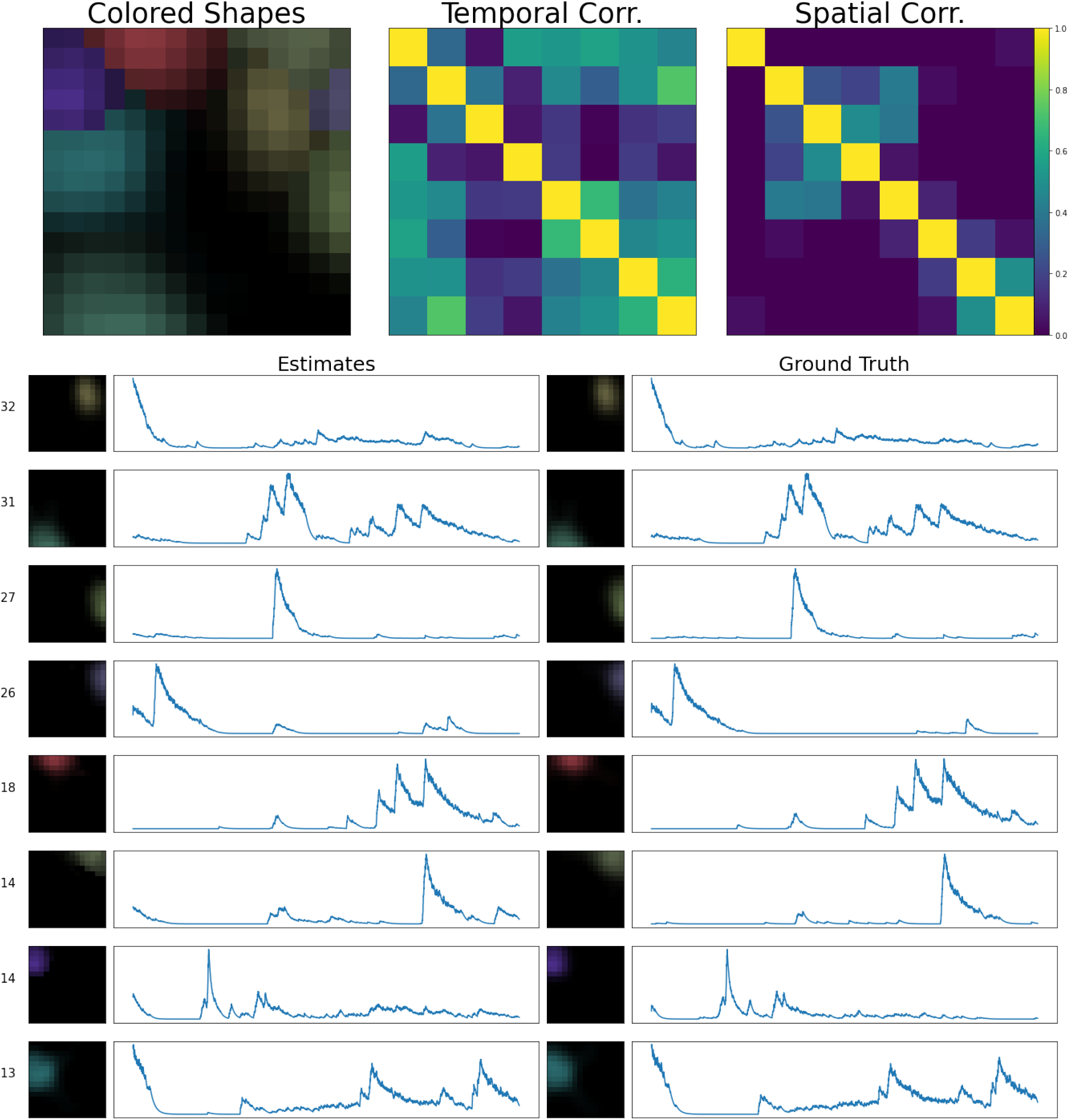
Sample frame of a simulated demixing video. Here, we show all components extracted from a spatial patch of the simulated dataset described in Fig. 6. In the lower half of this figure, we provide every estimated component’s spatial and temporal footprint in the left column. These estimates are ordered in terms of their maximum brightness (maximum of *a*_*i*_(*x*)*c*_*i*_(*t*)) in the video. Next to each component’s spatial footprint, we provide this maximum brightness value for reference. Each estimate is matched with its corresponding ground truth neural signal. In this example all of our estimated components match the ground truth with high accuracy. Each component is assigned a unique color (matching the colors assigned in Fig. 6). In the top left corner, we provide an aggregate image, showing a max-projection of each component (in its corresponding color) as it appears in the field of view. To the right of that panel, we provide temporal and spatial correlation matrices of our estimated components.

Next we quantify how our algorithm performs in different *R*_*c*_ regimes: how does accuracy break down as the density of cells per pixel increases? We compare several algorithms here: the proposed new algorithm (denoted “maskNMF”); the original localNMF algorithm from Buchanan et al. (2018); Suite2p from Pachitariu et al. (2016); and finally, an “oracle” algorithm that is provided the correct spatial component shapes and simply needs to estimate the temporal traces corresponding to these spatial components (via standard non-negative regression). This oracle algorithm is a useful “upper bound” for understanding signal recovery, because it tells us how accurately we can estimate the neurons’ temporal activities given *perfect* estimates of their spatial footprints.

To evaluate the above pipelines’ outputs on a simulated dataset, we compute two key metrics: recovery accuracy and false positive count (FPC). To calculate the recovery accuracy, we begin by matching the neuron estimates produced by a given pipeline with the ground truth. Specifically, for each ground-truth neuron, we identify a set of spatially similar estimated components, where similarity is measured via cosine similarity. Among those spatially similar components, we identify the most temporally similar one (again, temporal similarity is defined by cosine distance) *that has not yet been matched*. If no match is found by this process, the ground truth neuron is treated as unmatched, with temporal similarity 0. We then define the recovery accuracy as the average temporal similarity among all ground truth neurons and their respective matches. Then, we define the FPC as the number of estimates which were unmatched. Together, these two metrics give us a clear indication of whether a given pipeline is adequately demixing the signals in the video.

As expected, recovery accuracy deteriorates as the cell density *R*_*c*_ increases. However, our new approach significantly outperforms localNMF and Suite2p, both in terms of recovery accuracy and FPC; see Fig. 8. Furthermore, across many density values, our new approach performs near the level of the “oracle” algorithm. Therefore, the new pipeline both outperforms existing state-of-the-art pipelines and also achieves near optimal recovery accuracy over the dense simulated datasets analyzed here.

**Figure 8:**
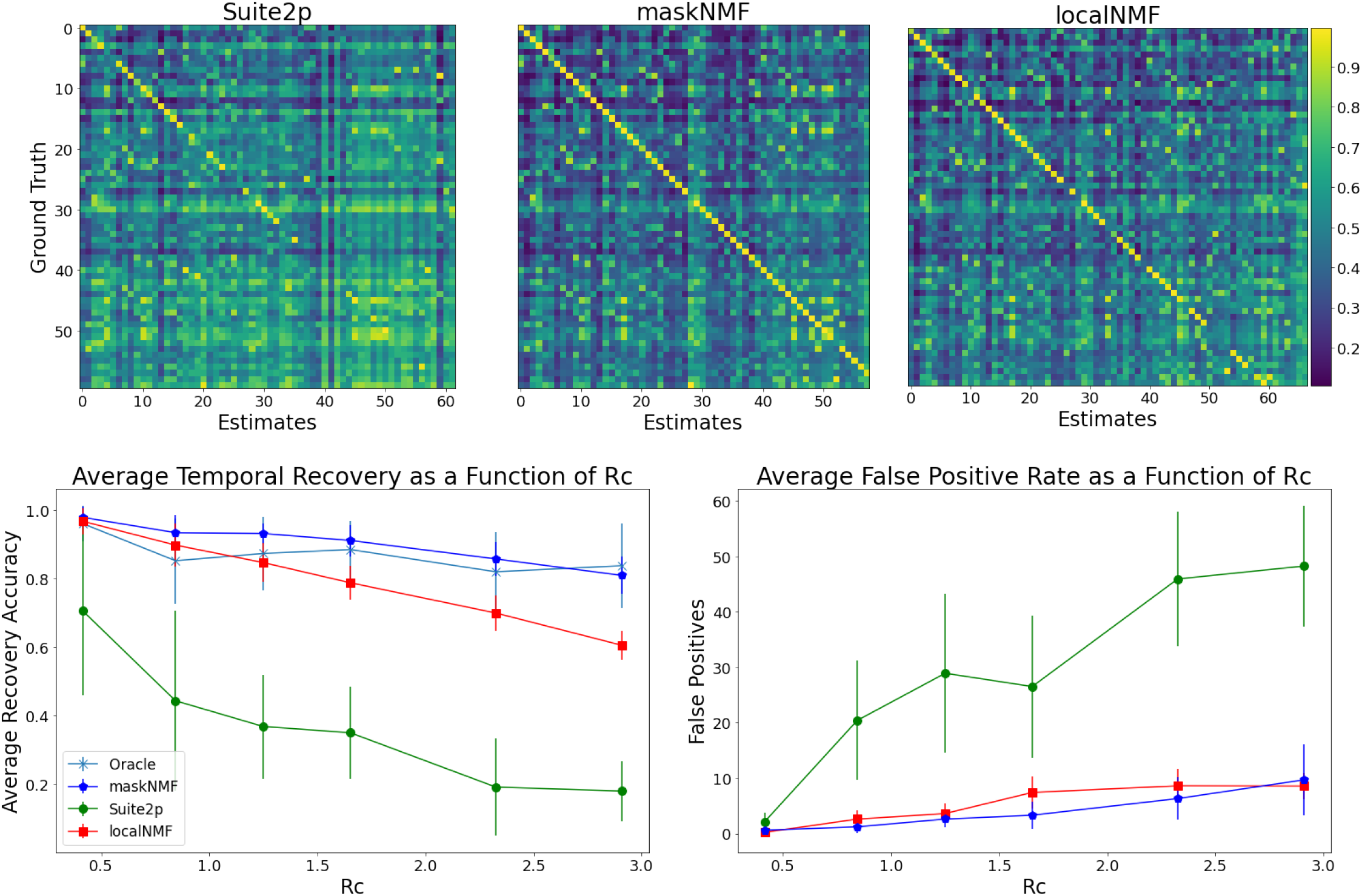
Summary of simulation comparisons. Top: Here we provide correlation matrices for several calcium imaging pipelines run on the simulated dataset from Fig. 6. For each of these correlation matrices, we match the pipelines’ estimates with the corresponding ground truth. The ground truth neurons are ordered by brightness in exactly the same way for each of these matrices. This allows us to visually compare the recovery of individual ground truth neurons across the pipelines. For the maskNMF pipeline, almost all entries are close to 1 across the diagonal, indicating that the pipeline’s estimates and ground truth neurons were matched accurately, and our algorithm has properly demixed these signals. For the localNMF pipeline, there are more entries along the diagonal of the matrix which are not close to 1, which indicates that the pipeline failed to fully recover those ground truth neurons. We ran the Suite2p pipeline in an automated fashion, choosing parameters to maximize performance at each density level (neuron overlap enabled, and neuropil estimation disabled, because our simulated videos did not contain any neuropil). Both localNMF and maskNMF significantly outperform Suite2p. **Bottom**: We provide a comparison of Suite2p, maskNMF, and localNMF using simulated datasets across a range of densities (*R*_*c*_). Errorbars indicate standard error over multiple replicates of the simulations. Note that Suite2p performs relatively poorly on dense data, while the new proposed pipeline tracks the best-case “Oracle” approach over the range of densities analyzed here.

### Application to real Bessel data

Now we turn to real Bessel imaging data. In this section, we examine a real Bessel imaging dataset whose estimated density is approximately *R*_*c*_ = 0.7. We start by examining the demixing video, which illustrates the various outputs of the pipeline (Fig. 9).

**Figure 9:**
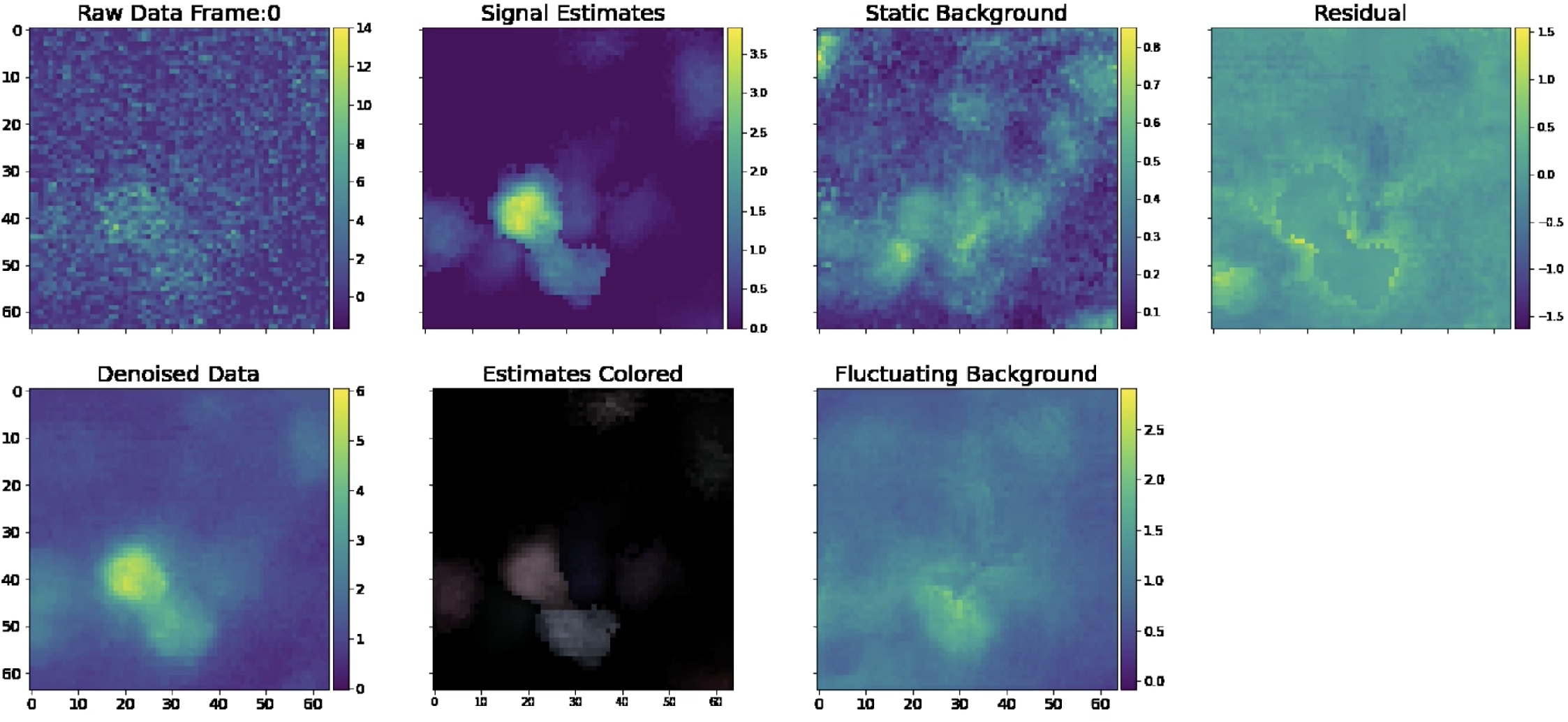
Single frame of a demixing video generated from real Bessel data. Similar to the frame provided in Fig. 6, we provide a frame of a demixing video on *real* data. Note that there is no Ground Truth panel here because this is a real dataset. Also note that because this is a real dataset, the algorithm estimates neuropil activity and the Fluctuating Background Frame is non-empty. We provide the full demixing video [here](https://drive.google.com/file/d/19GA7-LOxhT6FjacktyYDrKUb0fd0k6IR/view?usp=sharing).

To assess the quality of our signal estimates, we can visually compare these estimates to the activity from the denoised data. We can also examine the residual for any salient leftover signals. In the demixing video, there is very little leftover signal in the residual video, indicating that our estimates are of a high quality.

In Fig. 10, we provide a library of all components extracted from this field of view. Overall, demixing is successful here: no two neurons appear to be absorbing the same signal in the video. Furthermore, neurons that are spatially overlapping and temporally correlated are being successfully demixed. Thus the pipeline seems to work well on real Bessel data, in addition to the simulated results shown above.

**Figure 10:**
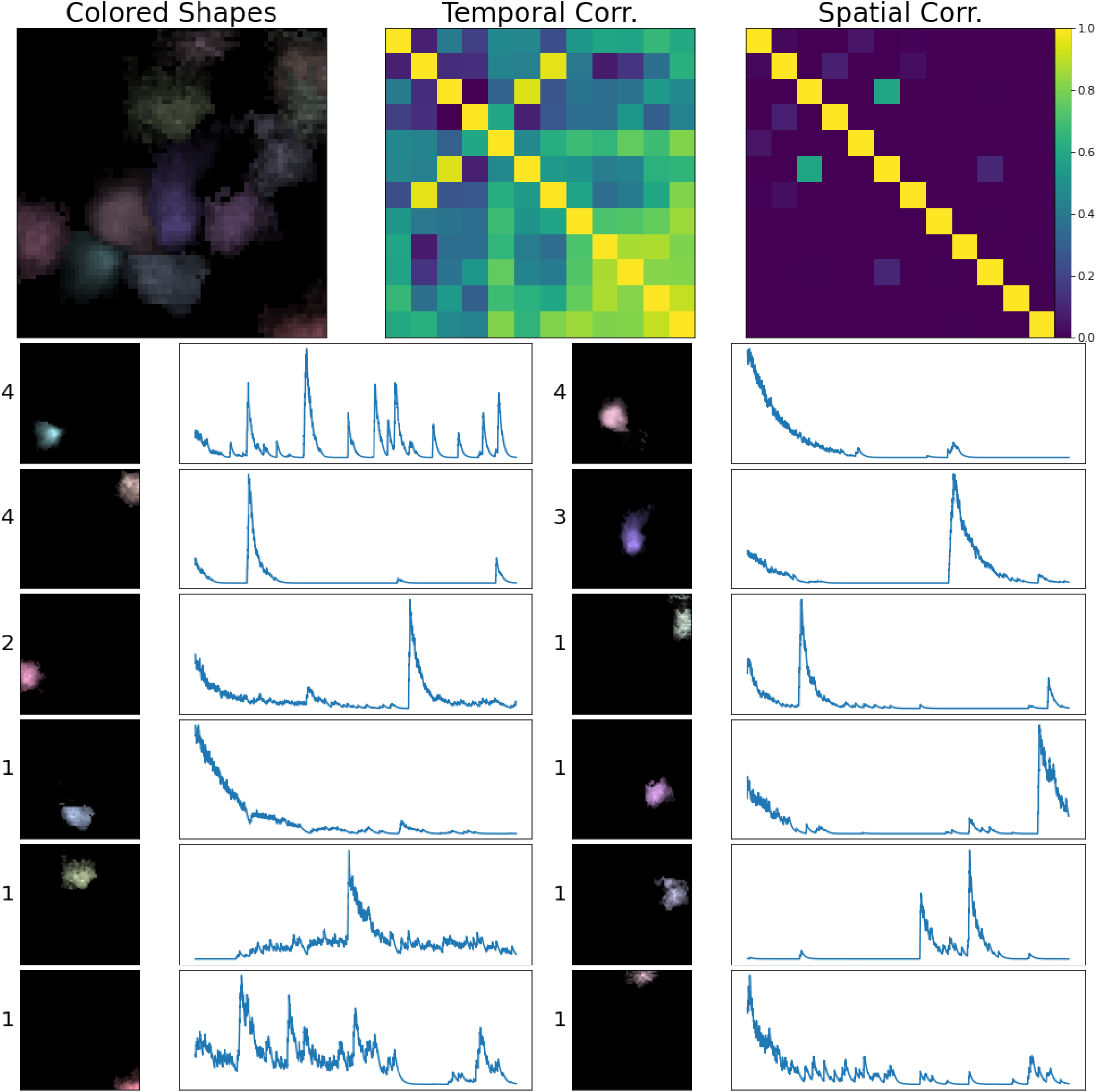
Real source extraction for Bessel data. Here, we show all components extracted from a spatial patch of the real Bessel imaging dataset, in a similar fashion to Fig. 7; the only difference here is that there is no ground truth. Each component is assigned a unique color (matching the colors assigned in Fig. 9).

### Application to real standard two-photon imaging data

To explore the flexibility of the proposed framework, we next evaluated our new pipeline on standard two-photon calcium imaging data. Here we show results on a two-photon dataset (one of the “standard” datasets used in (Giovannucci et al., 2019)) with estimated density *R*_*c*_ = 1.0. Our procedure for training the Mask R-CNN is identical to the one we used for Bessel imaging data, except we use a two-photon point spread function in the generative model (following (Zhou et al., 2020)) used to simulate spatial footprints.

In Fig. 11, we provide a single frame of the demixing video for this dataset. Close comparison of the signal estimates and the denoised data in the corresponding demixing video suggest that our algorithm is adequately identifying and demixing the large majority of signal present in this video. Similarly, close inspection of the residual indicate that there is very little significant leftover signal missed by the new pipeline.

**Figure 11:**
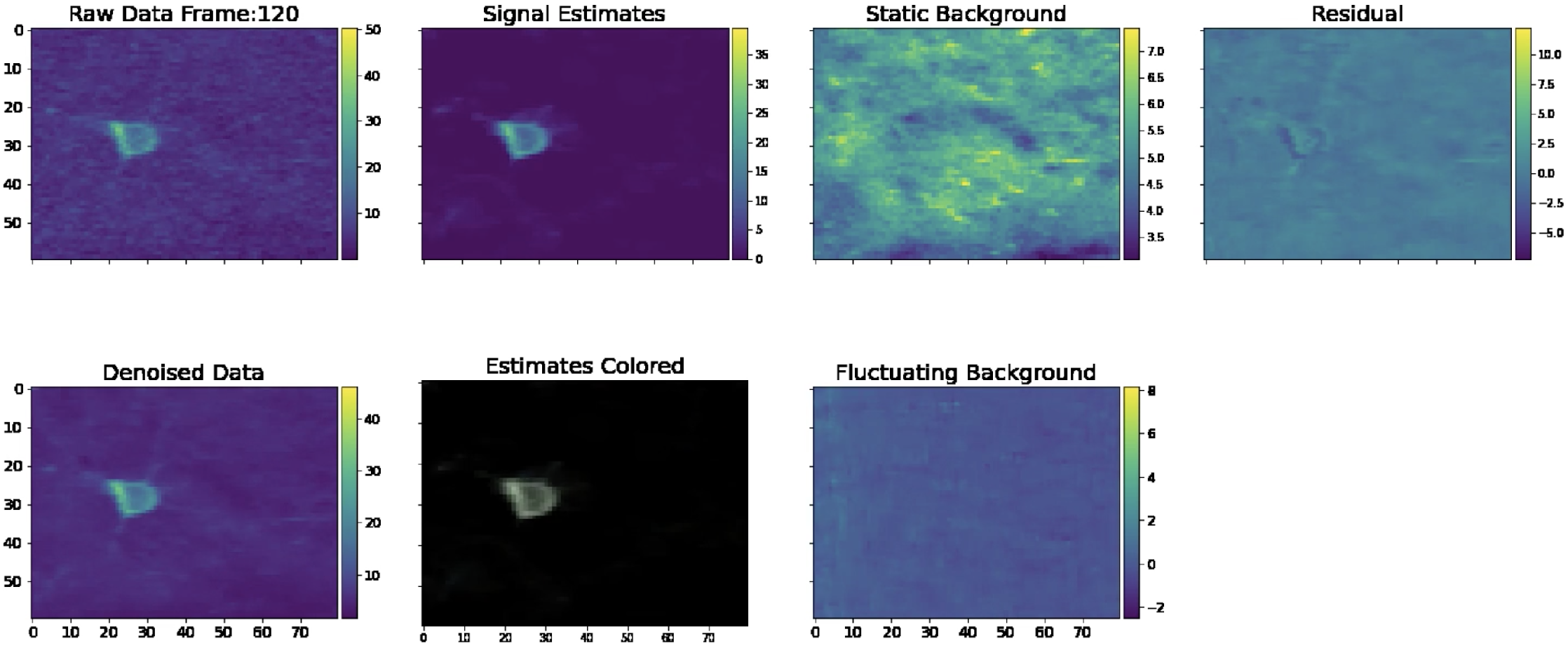
Single frame of a demixing video generated from real two-photon data. In this frame, we see the various components of the demixing movie, presented in the exact same manner as 9. The lack of visible signal in the mean-subtracted residual suggests that our algorithm has successfully extracted all signals. Unlike 9, we subtract the minimum value from each pixel of the “Signal Estimates” and “Estimated Colored” frame to allow the reader to more clearly see the most active cells in each frame. Note that this dataset contains both somatic and dendritic signals; our algorithm is able to adequately identify both. We provide a demixing video [here](https://drive.google.com/file/d/1Ew93O5x8ymVXmOscVsjn1CCBDHfkzwMn/view?usp=sharing).

Fig. 12 provides the library of neurons extracted over a spatial sub-patch of the field of view shown in Fig. 11. The estimated components largely correspond to full shapes (either apical dendrites or somas), as desired. In summary, we conclude that the proposed pipeline performs well on standard two-photon data, in addition to the Bessel data analyzed above.

**Figure 12:**
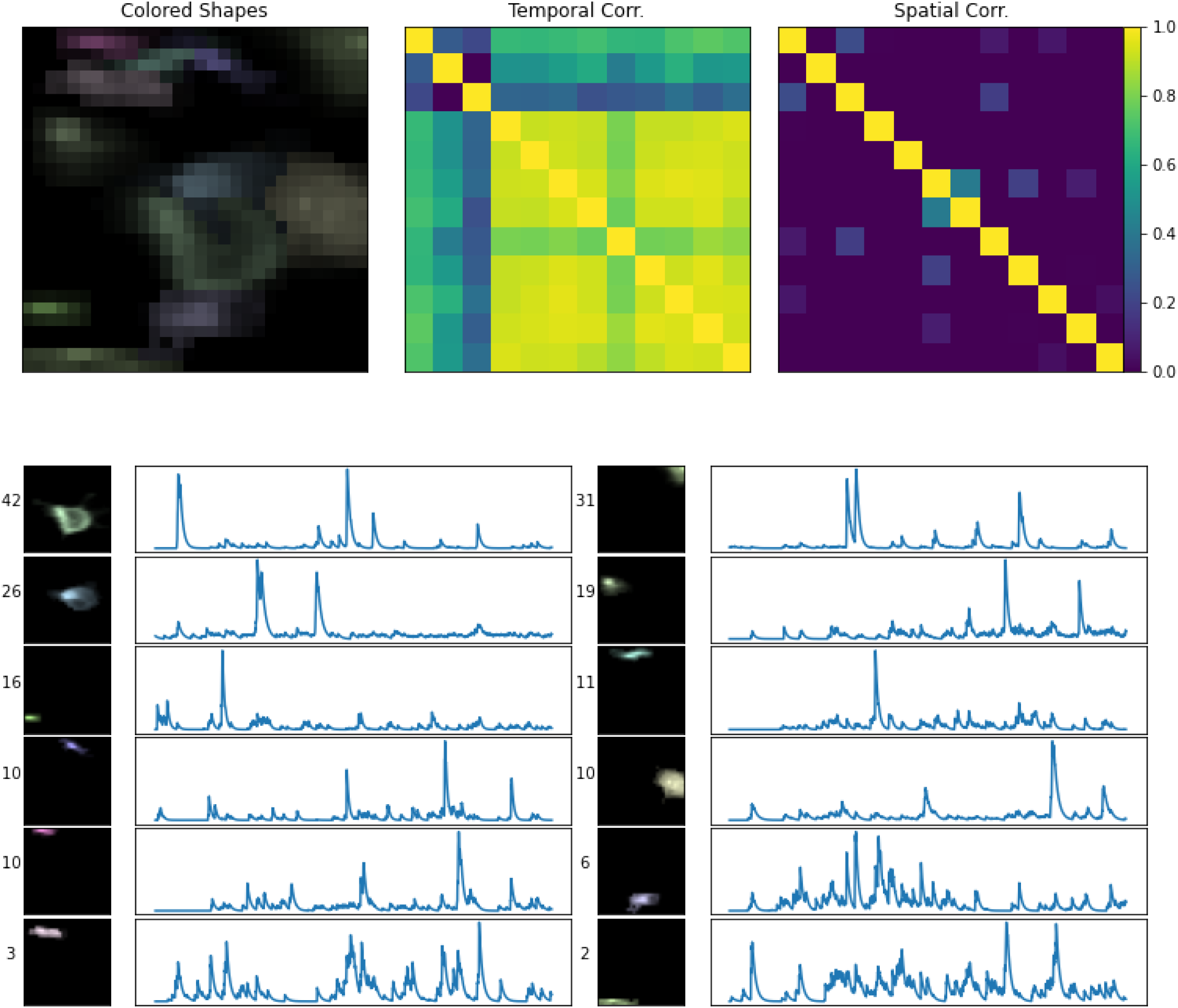
Real source extraction for two-photon data. Here, we show all components extracted from a subpatch of the real two-photon imaging dataset shown in Fig. 11. Conventions as in Fig. 10.

## Conclusion and future work

In this work, we have introduced an algorithm for calcium imaging analysis which can demix significantly denser data than was previously feasible. These computational advances are motivated by experimental progress in calcium imaging. In particular, methods such as Bessel imaging increase the number of cells that can be acquired with two-photon imaging, but at the cost of highly-mixed outputs of the imaging system.

We use a three-step approach to solve the demixing problem. First, we denoise the data. Then, we deconvolve the data (and denoise the output) to sparsen and better isolate neuron signals. Finally, we detect isolated neuron shapes using a neural network and ultimately demix the signals detected in the video data. The proposed pipeline is fully automated, requiring no manual data annotation; specifically, we use simulated neuron shapes based on electron microscopy data (not manually-annotated data) to train the detection neural network.

The proposed pipeline performs well on a diverse set of real and simulated datasets of varying cell densities. On simulated data, our new approach outperforms state of the art calcium imaging analysis pipelines, particularly in the densest regime. It also achieves near optimal demixing results at densities in which existing pipelines largely break down. Thus the new pipeline opens the door to more densely-mixed experiments targeting simultaneous imaging of significantly larger neural populations than is feasible with previous demixing pipelines.

Our work leaves open many interesting open questions. First, we used a standard object detection architecture (Mask R-CNN) from the computer vision community; perhaps a more specialized architecture might provide stronger performance. Another interesting possibility would be to use a neural network which can take advantage of slightly more temporal context. For example, instead of taking as input a single frame and detecting objects, a more sophisticated network might look at a 5 − 10 contiguous frames and use the spatiotemporal context to identify neurons; see e.g. (Soltanian-Zadeh et al., 2019) for a related approach. Another important challenge in calcium imaging analysis is the task of discarding bad signal estimates, either as a post-processing step or during the demixing iterations. Using our framework for simulating data, we could build a model which can discriminate between shapes which look like calcium footprints and those that do not; this may lead to further performance gains. See (Pachitariu et al., 2016; Giovannucci et al., 2019) for related manually-trained approaches. Finally, the speed of our methods and structure of our software tools opens the up the possibility for numerous interesting online analyses on GPUs; see (Cai et al., 2023) for an existing approach in this direction.

## Acknowledgements

We would like to thank Marcus Triplett, Ben Antin, Kushal Kolar, Anna Yoon, and Abhishek Shah for helpful conversations. This work was supported by NIH 1U01NS103489-01, NSF Neuronex DBI-1707398, NIH 5U19NS104649, and NIH 5U19NS107613.

## A. Appendix: Superpixel initialization details

Buchanan et al. (2018) introduce a correlation based method (called “superpixels”) for identifying neural signals. In this paper, we incorporate the superpixels initialization procedure in our multi-pass approach for demixing neural signals. We first initialize a set of neurons using maskNMF, and run the localNMF demixing method accordingly. We then examine the “residual movie” (the movie obtained by subtracting away all estimated signals) and use “superpixels” initialization to identify any remaining signals. This procedure can be iterated if desired.

To understand the superpixel method, consider a graph, *G*, whose vertices are the pixels from the video field of view. We define two vertices (pixels) to be connected by an edge if and only if the pixels which they represent are adjacent and have a correlation that is above a certain correlation threshold. Large connected components in the graph *G* describe a cluster of highly correlated pixels in the imaging video, and likely represent neural signals. Buchanan et al. (2018) proposed the following method for calculating the correlation between two pixels,

*x* ∈ *R*^*T*^ and *y* ∈ *R*^*T*^. The first step serves as a lossy denoiser: we threshold the pixels based on their median absolute deviation (MAD):

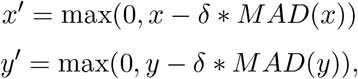

where *δ* is a nonnegative integer. In practice, we usually set *δ* = 2. Then, the correlation between *x* and *y* is defined to be the Pearson Correlation Coefficient between *x*′ and *y*′:

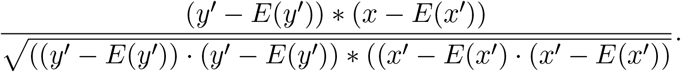

We note that this correlation metric makes the overall superpixel procedure sensitive to its correlation threshold. This is further exacerbated by the fact that PMD denoising empirically increases the correlation between pixels which do not contain any isolated neural activity (since background neuropil signal is retained while noise is decreased). To remedy this, we modify the correlation metric: we calculate the Pearson Correlation Coefficient between *x*′ + *n*_1_ and *y*′ + *n*_2_, where *n*_1_ and *n*_2_ are independent and identically distributed Gaussian noise vectors, drawn from a distribution with mean 0 and variance 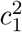. We interpret 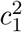 as a crude “leftover” post-compression noise term; we have found that values around 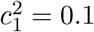 are effective. The resulting correlation becomes:

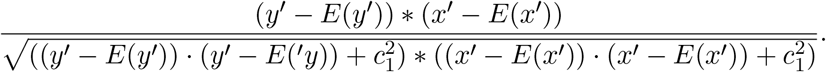

This serves to decrease the correlation values between dim pixels while leaving correlations between bright pixels relatively unchanged, improving the overall robustness of the superpixel approach, and decreasing its dependence on the precise value of the correlation threshold.

### B Overview and notation for localNMF

Here, we outline the localNMF algorithm, with an emphasis on the ways we have modified the original methods presented by the authors of (Buchanan et al., 2018). The key insight to all implementations here are that we can leverage the sparsity of *U* and low rank of *U* and *V* to achieve computationally faster and more memory-efficient demixing. Furthermore, the use of *U* and *V*, instead of the expanded movie 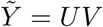, allows us to move the compressed data onto GPUs (which typically have very limited memory), leading to significant additional acceleration.

First, we introduce some notation and basic facts:

1. The imaging video, *Y*, has *d* pixels and *T* frames. It is a *d* x *T* matrix.
2. When we perform a HALS spatial or temporal regression update, we assume that we have *N* total neurons.
3. The PMD decomposition, 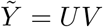 has rank *K*; that is, *U* is a *d* x *K* matrix, and *V* is a *K* x *T* matrix. *U* is a block-sparse matrix.
4. We have a spatial matrix, *A*, which has dimensions *d* x *N*. This matrix describes the spatial footprints of all neurons. It is sparse because neural signals are spatially local.
5. We have a temporal matrix, *C*, which has dimensions *T* x *k*. This matrix describes the spatial footprints of all neural signals.
6. As in the main text, we let *B* denote the *d* x *T* matrix describing the overall estimate of the background in our NMF model. *B* is the sum of two terms, a static, baseline background (*B*_*s*_) and a fluctuating background (*B*_*f*_). The *d* × 1 vector, *b*, describes the value of the static background *B*_*s*_ at every pixel.
7. *W* is a sparse matrix, which we call the “ring matrix.” It is used in the modeling of *B*_*f*_ ; see (Zhou et al., 2018) and below for more details. It has dimensions *d* x *d* and is used to estimate the fluctuating background.
8. During the localNMF procedure, we keep track of a sparse *d* × *N* binary matrix, *S*, whose *i*-th column describes the spatial support of the *i*-th neural signal in *A*.

#### Algorithm 1

RingLocalNMF

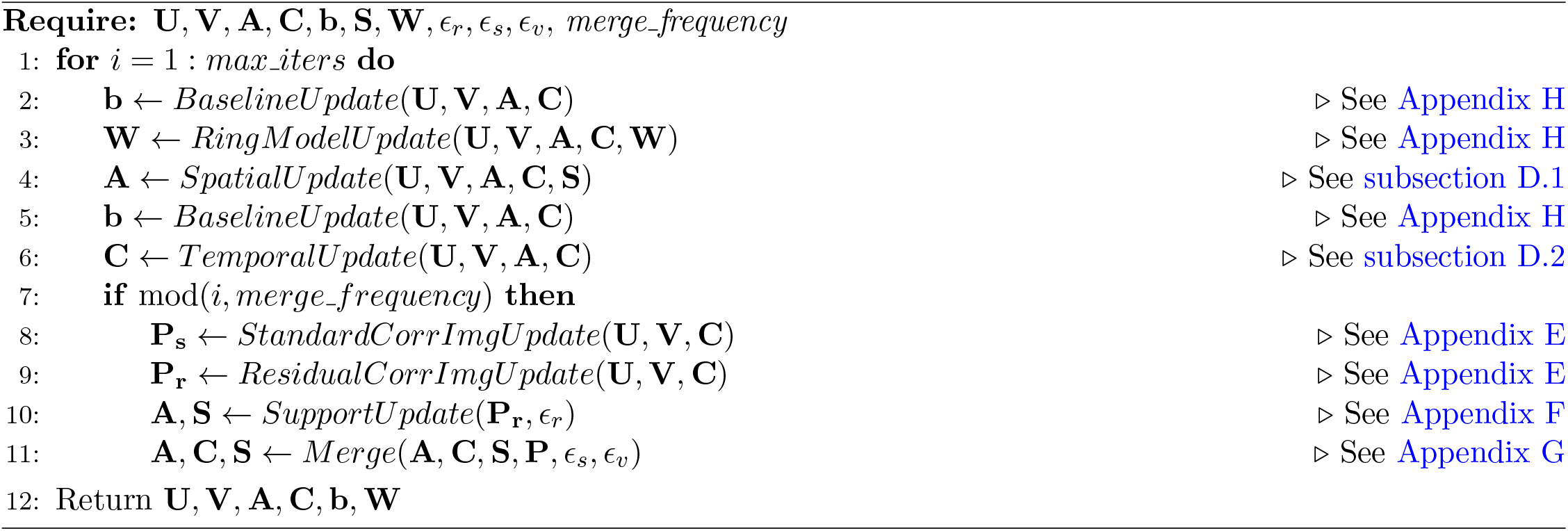

### C. Auxiliary low-rank factorizations

Here, we describe some key factorizations which we will use repeatedly in the following sections to accelerate the computation.

First, as described above, we will frequently use the original PMD compressed representation 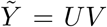 times, we will find it computationally useful to use an orthonormal basis for *V* ; therefore, at the start of this method, we run a QR factorization on *V* to arrive at the factorization 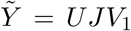 where *V*_1_ is an orthogonal matrix.

We also often refer to a so-called “residual” matrix:

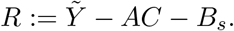

We want to express all terms in the decomposition in the temporal *V* basis. To do so, we first express *C* in terms of the *V* basis,

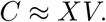

Here *X* is an *N* × *K* matrix, which we can efficiently find with a standard least squares solver. Next, we factorize the static baseline *B*_*s*_ as *B*_*s*_ = *b***1**_*T*_. We can express **1**_*T*_ using the *V* basis as follows: **1**_*T*_ = *sV*. To ensure that such a vector, *s*, exists, we append a vector of 1’s to *V* at the start of the localNMF algorithm guaranteeing that the vector of 1’s belongs to the rowspace of *V* .

Putting these individual factorizations together, we arrive at the following expression for the residual matrix:

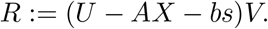

We can arrive at a similar type of expression if we use the orthogonal basis, *V*_1_:

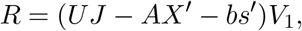

where, of course, *X*^’^ and *s*^’^ differ from *X* and *s* because they must be estimated using the appropriate temporal basis matrix (*V*_1_).

Finally, we express the fluctuating background in terms of the *V*_1_ temporal basis. As described in (Zhou et al., 2018), the ring model is defined as

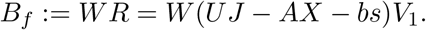

### D HALS update steps

In this section, we describe how to conduct temporal and spatial update steps in localNMF.

#### D.1 Spatial HALS

We begin by describing the procedure for performing spatial HALS updates. For each neuron, *i*, we perform the following update:

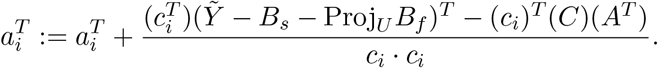

Because this step updates all of the *d* pixels of 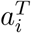 we next set all pixels which lie outside of the support of *a*_*i*_ (given by the *i*-th column of *S*) to 0. Note that we project the fluctuating background term, *B*_*f*_, onto the *U* subspace; this step helps to denoise and regularize the fluctuating background estimate.

Using the factorizations described above, we have the following update rule in terms of *U* and *V* :

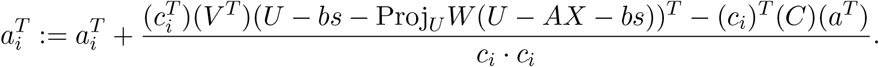

If we precompute the appropriate quantities across HALS iterations and then conduct the remaining matrix multiplications in the appropriate order (ensuring that the result of every matrix multiplication “collapses” to a single vector instead of expanding into a large matrix), we can compute all spatial updates much more efficiently than previously possible. The asymptotic runtime for a full set of HALS updates (one for each of the *N* neural signals) using the *U* and *V* representation is *O*(*NK*(*d* + *T*)). On the other hand, the asymptotic runtime of running HALS on the full video 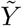 would be *O*(*NdT*); recall that typically *K* will be much smaller than *d* or *T*. Note that this analysis does not account for the sparsity of *U* and *A*, which leads to further speedups.

#### D.2 Temporal HALS

We can use similar methods as above to efficiently calculate temporal HALS updates. During the temporal HALS updates, for each neural signal, *i*, we update *c*_*i*_ as follows:

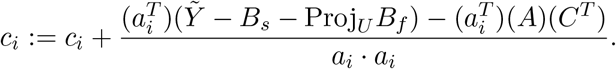

As we did in the spatial HALS updates, we can express the above update using the compressed *U* and *V* representations,

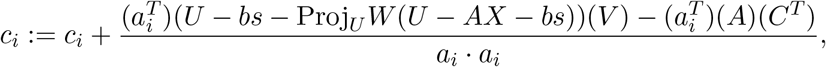

nd gain similar speedups (*O*(*NK*(*d* + *T*)) time rather than the original *O*(*NdT*) cost per iteration on the full matrix 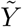). Note that like the spatial HALS update step, we also project *B*_*f*_ onto the *U* subspace here.

#### E. Correlation image calculations

In our enhanced implementation of the localNMF pipeline, we use two correlation images. The first is the “standard” correlation image described in (Buchanan et al., 2018), which gives the correlation between every column of *C* and every pixel of 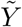 The second is a new “residual correlation image,” which we use for updating the spatial supports of all neural signals. We now show how to efficiently compute the residual correlation image for each neural signal by exploiting the orthogonalized PMD representation, 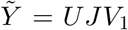 We also note that the same overall approach can be used to arrive at the standard correlation image.

To compute the residual correlation image for a neural component, given by a spatial fluorescence profile *a*_*i*_ and temporal profile *c*_*i*_, we first subtract all other signals (aside from *a*_*i*_ and *c*_*i*_) from the denoised movie, 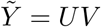

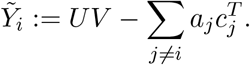

Then the *i*-th residual correlation image is defined as the standard correlation image between *c*_*i*_ and 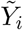

We begin by normalizing the columns of the matrix *C*. Specifically, for each column of *C*, we first center the column vector and then divide this result by its norm; this results in 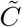 a *T* × *N* matrix. We let 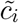 refer to the *i*-th normalized neuron trace (it is the *i*-th column of 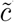).

We approximate *C* as a product *X*′*V*_1_, by least squares. Then, *AC* can be approximated by *AX*′*V*_1_, and the centered residual movie can then be written in the following factorized form:

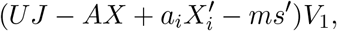

where *m* denotes the *d* × 1 vector describing the mean of every pixel in 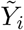

To complete the correlation calculation, we need to find the norm of the centered movie. We start by finding the square of the pixelwise norm for this centered residual movie:

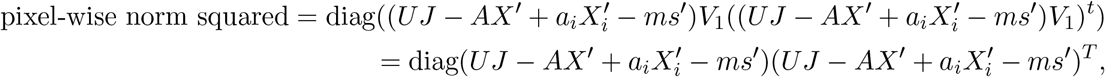

where the *V*_1_ term disappears because it has orthogonal rows. Note that we can complete this calculation by manipulating *d* x *K* matrices (instead of the significantly larger *d* x *T* matrices with which we started). The above computation is efficient, under two assumptions. First, as above, we must compute matrix products in the appropriate order (matrix products should “collapse” to lower dimensional matrices, not expand). Second, we can also precompute the diagonal of many of the above quantities; this results in further runtime savings since we repeatedly compute the residual correlation image.

We take the square root of the above result to get the pixel-wise norm of the residual movie. Finally, to estimate the residual correlation image for neuron *i*, we multiply:

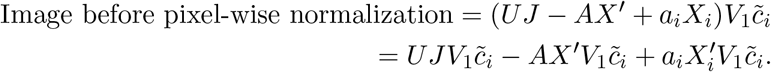

As usual, we take care in conducting these matrix operations in the right order to ensure that we are only manipulating matrix-vector products. The final result is a *d* × 1 unnormalized correlation image. We divide every row of this result by the corresponding pixelwise norm (computed above) to get our final result.

#### F Neuron Support Updates

We briefly note that in (Buchanan et al., 2018), the LocalNMF algorithm received as input a correlation threshold, *E*_*r*_. At every iteration, the support of neural signal *i* was computed by first thresholding every neuron’s *standard* correlation image above the value *E*_*r*_, and considering the connected region between this thresholded image and the neuron’s current spatial support.

This approach has a significant drawback – that a single correlation image may not adequately update every neural signal’s support adequately. To remedy this, we introduce two changes. First, instead of using the standard correlation image, we use the residual correlation image. This is particularly important in the case of dense projective imaging data, where signals exhibit a large degree of spatial overlap. Second, the threshold parameter *ϵ*_*r*_ is no longer a hard threshold; instead, we take the maximum value of each neuron’s residual correlation image and multiply it by 0 *< ϵ*_*r*_ *<* 1. This produces a fine-tuned correlation threshold for each individual correlation image, leading to more robust results.

#### G Neuron Merge Updates

In exactly the same fashion as in (Buchanan et al., 2018), we use the standard correlation images of the neurons of *A* to decide which neural signals should be merged. This approach takes as input two “threshold” parameters, 0 *< ϵ*_*t*_ *<* 1 and 0 *< ϵ*_*v*_ *<* 1. We first truncate all of the standard correlation images of *A* with the threshold *ϵ*_*t*_. Then, for every possible pair of neural signals, we compute the degree of overlap between their truncated correlation images. This degree is expressed as a fraction of the total number of pixels in their supports and is therefore nonnegative and at most 1.

Conceptually, this computation defines a graph *G*, whose vertices consist of each of the neural signals in *A*. In this graph, we define two vertices to be connected by an edge if and only if their degree of overlap exceeds the threshold *ϵ*_*v*_. We then identify all of the connected components in *G*. Each connected component is defined by a set of vertices, *M*. We merge the neural signals corresponding to the vertices of *M*. Computationally, for each connected component *M*, we sum the signals,

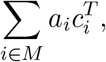

We then perform a rank-1 NMF on this result to identify a single, merged spatial and temporal signal. We define the support of this resultant signal to be the union of the supports of the neural signals that were merged.

#### H. Background Updates

Recall that we model the background, *B*, as a sum of a static baseline fluorescence term, *B*_*s*_ and a time-varying term, *B*_*f*_ :

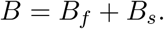

We estimate *B*_*s*_ during each HALS iteration; it is defined as the difference between the pixelwise mean of *UV* and *AC*^*T*^ .

To estimate *B*_*f*_, we adapt the ring model from CNMF-E (Zhou et al., 2018). This approach seeks to model the fluctuating background as:

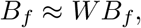

where *W* is a linear operator which attempts to explain the activity of every pixel of *B*_*f*_ exclusively as a linear combination of the pixels of *B*_*f*_ which lie on a ring of radius *r* from the center pixel. In practice, one can fit this model with the following simplification: instead, we seek a *W* which minimizes

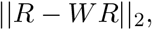

where *R* := *UV* − *AC*^*T*^ − *B*_*s*_, and *W* is a highly sparse *d* x *d* ring matrix.

In this paper, we further simplify the ring model with two restrictions. First, we require that for every row of *W*, all nonzero entries have the same value; this is the so-called constant ring assumption. For each pixel, we call this constant value the “ring weight” for that pixel. Second, if the *i*-th pixel belongs to the support of *A*, then the *i*-th column of *W* consists of all zeros; in other words, no ring can contain pixels belonging to the support of *A*.

A major benefit of the constant assumption is that it allows for a significantly faster model fit; we only need to update a single parameter per pixel (the ring weight). Using the compressed representation 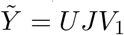 we can express this objective function as:

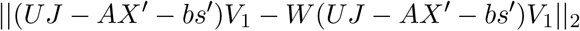

For any given pixel, *i*, let *x* denote the sum of the pixels belonging to its ring, and let *y* denote the *i*-th pixel of *R*. To find the ring weight in this scenario, it suffices to minimize:

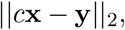

which, via standard least squares regression, is:

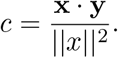

Viewed in this light, we can compute the updated weights for every pixel as a series of matrix operations. Let *W*_0_ denote the ring matrix whose nonzero elements all have value 1 (i.e. every ring weight is 1). We can express the 1 x *T* vector of weights, *w*, which solves the above optimization problem, as follows:

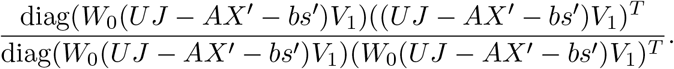

Here, division is an element-wise division between the vector in the denominator and the vector in the numerator. As above, in this expression, all *V*_1_ terms cancel due to orthogonality, which yields the following:

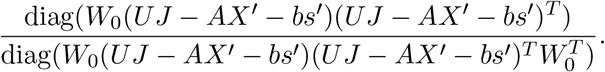

This expression is now significantly easier to compute: we are manipulating matrices of size at most *d* x *K* now (instead of the much larger *d* x *T*).

#### I Open source implementations

Here, we provide a list containing the open source implementations of all code used in the maskNMF pipeline:

1. Accelerated NoRMCorre: [JNoRMCorre](https://github.com/apasarkar/jnormcorre).
2. Accelerated PMD Compression and Denoising: [PMD](https://github.com/apasarkar/localmd).
3. maskNMF: [maskNMF](https://github.com/apasarkar/masknmf).
4. localNMF demixing: [rlocalnmf](https://github.com/apasarkar/rlocalnmf).
5. Dockerized Dash app / platform: [App](https://github.com/apasarkar/masknmf_full_pipeline) and [App Docs](https://masknmf-full-pipeline.readthedocs.io/en/latest/).
6. Napari plugin for high frame rate local data visualization: [plugin](https://github.com/apasarkar/napari-masknmf).

## References

Apthorpe, N. J., Riordan, A. J., Aguilar, R. E., Homann, J., Gu, Y., Tank, D. W., and Seung, H. S. (2016). Automatic neuron detection in calcium imaging data using convolutional networks. In Proceedings of the 30th International Conference on Neural Information Processing Systems, NIPS’16, page 3278–3286, Red Hook, NY, USA. Curran Associates Inc.

Arora, S., Ge, R., Kannan, R., and Moitra, A. (2016). Computing a nonnegative matrix factorization -provably. SIAM J. Comput., 45(4):1582–1611.

Bao, Y., Soltanian-Zadeh, S., Farsiu, S., and Gong, Y. (2021). Segmentation of neurons from fluorescence calcium recordings beyond real-time. Nat Mach Intell, 3(7):590–600.

Batson, J. and Royer, L. (2019). Noise2self: Blind denoising by self-supervision.

Berens, P., Freeman, J., Deneux, T., Chenkov, N., McColgan, T., Speiser, A., Macke, J. H., Turaga, S. C., Mineault, P., Rupprecht, P., Gerhard, S., Friedrich, R. W., Friedrich, J., Paninski, L., Pachitariu, M., Harris, K. D., Bolte, B., Machado, T. A., Ringach, D., Stone, J., Rogerson, L. E., Sofroniew, N. J., Reimer, J., Froudarakis, E., Euler, T., Román Rosán, M., Theis, L., Tolias, A. S., and Bethge, M. (2018). Community-based benchmarking improves spike rate inference from two-photon calcium imaging data. PLoS Comput. Biol., 14(5):e1006157.

Buchanan, E. K., Kinsella, I., Zhou, D., Zhu, R., Zhou, P., Gerhard, F., Ferrante, J., Ma, Y., Kim, S., Shaik, M., Liang, Y., Lu, R., Reimer, J., Fahey, P., Muhammad, T., Dempsey, G., Hillman, E., Ji, N., Tolias, A., and Paninski, L. (2018). Penalized matrix decomposition for denoising, compression, and improved demixing of functional imaging data. bioRxiv, page 334706.

Cai, C., Dong, C., Friedrich, J., Rozsa, M., Pnevmatikakis, E. A., and Giovannucci, A. (2023). FIOLA: an accelerated pipeline for fluorescence imaging online analysis. Nat. Methods, 20(9):1417–1425.

Cai, C., Friedrich, J., Singh, A., Eybposh, M. H., Pnevmatikakis, E. A., Podgorski, K., and Giovannucci, A. (2021). Volpy: Automated and scalable analysis pipelines for voltage imaging datasets. PLOS Computational Biology, 17(4):1–27.

Charles, A. S., Cermak, N., Affan, R. O., Scott, B. B., Schiller, J., and Mishne, G. (2022). GraFT: Graph filtered temporal dictionary learning for functional neural imaging. IEEE Trans Image Process, 31:3509–3524.

Charles, A. S., Song, A., Gauthier, J. L., Pillow, J. W., and Tank, D. W. (2019). Neural Anatomy and Optical Microscopy (NAOMi) Simulation for evaluating calcium imaging methods. bioRxiv, page 726174.

Cichocki, A., Zdunek, R., and Amari, S.-i. (2007). Hierarchical ALS algorithms for nonnegative matrix and 3D tensor factorization. In International Conference on Independent Component Analysis and Signal Separation, pages 169–176. Springer.

Denis, J., Dard, R. F., Quiroli, E., Cossart, R., and Picardo, M. A. (2020). Deepcinac: A deep-learning-based python toolbox for inferring calcium imaging neuronal activity based on movie visualization. eNeuro, 7(4).

Dolev, N., Pinkus, L., and Rivlin-Etzion, M. (2019). Segment2p: Parameter-free automated segmentation of cellular fluorescent signals. bioRxiv, 832188.

Ferran, D. and Hamprecht, F. A. (2014). Sparse space-time deconvolution for calcium image analysis. In Proceedings of the 27th International Conference on Neural Information Processing Systems -Volume 1, NIPS’14, page 64–72, Cambridge, MA, USA. MIT Press.

Friedrich, J., Yang, W., Soudry, D., Mu, Y., Ahrens, M. B., Yuste, R., Peterka, D. S., and Paninski, L. (2017a). Multi-scale approaches for high-speed imaging and analysis of large neural populations. PLoS computational biology, 13(8):e1005685.

Friedrich, J., Zhou, P., and Paninski, L. (2017b). Fast online deconvolution of calcium imaging data. PLOS Computational Biology, 13(3):e1005423.

Frostig, R., Johnson, M., and Leary, C. (2018). Compiling machine learning programs via high-level tracing.

Giovannucci, A., Friedrich, J., Gunn, P., Kalfon, J., Brown, B. L., Koay, S. A., Taxidis, J., Najafi, F., Gauthier, L., Zhou, P., Khakh, B. S., Tank, D. W., Chklovskii, D. B., and Pnevmatikakis, E. A. (2019). CaImAn an open source tool for scalable calcium imaging data analysis. eLife, 8.

Halko, N., Martinsson, P. G., and Tropp, J. A. (2011). Finding structure with randomness: Probabilistic algorithms for constructing approximate matrix decompositions. SIAM Review, 53(2):217–288.

He, K., Gkioxari, G., Dollár, P., and Girshick, R. (2017). Mask R-CNN. In 2017 IEEE International Conference on Computer Vision (ICCV), pages 2980–2988.

Inan, H., Erdogdu, M. A., and Schnitzer, M. (2017). Robust estimation of neural signals in calcium imaging. In Guyon, I., Luxburg, U. V., Bengio, S., Wallach, H., Fergus, R., Vishwanathan, S., and Garnett, R., editors, Advances in Neural Information Processing Systems, volume 30. Curran Associates, Inc.

Jewell, S. and Witten, D. (2018). Exact spike train inference via (0) optimization. The annals of applied statistics, 12(4):2457–2482.

Jewell, S. W., Hocking, T. D., Fearnhead, P., and Witten, D. M. (2019). Fast nonconvex deconvolution of calcium imaging data. Biostatistics, 21(4):709–726.

Ji, N., Freeman, J., and Smith, S. L. (2016). Technologies for imaging neural activity in large volumes. Nature Neuroscience, 19(9):1154–1164.

Kaifosh, P., Zaremba, J. D., Danielson, N. B., and Losonczy, A. (2014). Sima: Python software for analysis of dynamic fluorescence imaging data. Frontiers in Neuroinformatics, 8:80.

Kirschbaum, E., Bailoni, A., and Hamprecht, F. A. (2020). Disco: Deep learning, instance segmentation, and correlations for cell segmentation in calcium imaging. In Martel, A. L., Abolmaesumi, P., Stoyanov, D., Mateus, D., Zuluaga, M. A., Zhou, S. K., Racoceanu, D., and Joskowicz, L., editors, Medical Image Computing and Computer Assisted Intervention –MICCAI 2020, pages 151–162, Cham. Springer International Publishing.

Klibisz, A., Rose, D., Eicholtz, M., Blundon, J., and Zakharenko, S. (2017). Fast, simple calcium imaging segmentation with fully convolutional networks. In Cardoso, M. J., Arbel, T., Carneiro, G., Syeda-Mahmood, T., Tavares, J. M. R. S., Moradi, M., Bradley, A., Greenspan, H., Papa, J. P., Madabhushi, A., Nascimento, J. C., Cardoso, J. S., Belagiannis, V., and Lu, Z., editors, Deep Learning in Medical Image Analysis and Multimodal Learning for Clinical Decision Support, pages 285–293, Cham. Springer International Publishing.

Krull, A., Buchholz, T.-O., and Jug, F. (2018). Noise2void -learning denoising from single noisy images.

Lecoq, J., Oliver, M., Siegle, J. H., Orlova, N., Ledochowitsch, P., and Koch, C. (2021). Removing independent noise in systems neuroscience data using deepinterpolation. Nature Methods, 18(11):1401–1408.

Lehtinen, J., Munkberg, J., Hasselgren, J., Laine, S., Karras, T., Aittala, M., and Aila, T. (2018). Noise2noise: Learning image restoration without clean data.

Lu, R., Sun, W., Liang, Y., Kerlin, A., Bierfeld, J., Seelig, J. D., Wilson, D. E., Scholl, B., Mohar, B., Tanimoto, M., Koyama, M., Fitzpatrick, D., Orger, M. B., and Ji, N. (2017). Video-rate volumetric functional imaging of the brain at synaptic resolution. Nature Neuroscience, 20(4):620–628.

Maruyama, R., Maeda, K., Moroda, H., Kato, I., Inoue, M., Miyakawa, H., and Aonishi, T. (2014). Detecting cells using non-negative matrix factorization on calcium imaging data. Neural Networks, 55:11 –19.

Mukamel, E. A., Nimmerjahn, A., and Schnitzer, M. J. (2009). Automated Analysis of Cellular Signals from Large-Scale Calcium Imaging Data. Neuron, 63(6):747–760.

Napari (2021). napari: a multi-dimensional image viewer for python.

Pachitariu, M., Packer, A., Pettit, N., Dagleish, H., Hausser, M., and Sahani, M. (2013). Extracting regions of interest from biological images with convolutional sparse block coding. In Proceedings of the 26th International Conference on Neural Information Processing Systems - Volume 2, NIPS’13, page 1745–1753, Red Hook, NY, USA. Curran Associates Inc.

Pachitariu, M., Stringer, C., Schröder, S., Dipoppa, M., Rossi, L. F., Carandini, M., and Harris, K. D. (2016). Suite2p: Beyond 10,000 neurons with standard two-photon microscopy. bioRxiv, pages 061507–061507.

Petersen, A., Simon, N., and Witten, D. (2018). SCALPEL: Extracting neurons from calcium imaging data. The Annals of Applied Statistics, 12(4):2430–2456.

Pnevmatikakis, E. A. and Giovannucci, A. (2017). Normcorre: An online algorithm for piecewise rigid motion correction of calcium imaging data. Journal of Neuroscience Methods, 291:83–94.

Pnevmatikakis, E. A. and Paninski, L. (2013). Sparse nonnegative deconvolution for compressive calcium imaging: algorithms and phase transitions. In Burges, C. J. C., Bottou, L., Welling, M., Ghahramani, Z., and Weinberger, Q., editors, Advances in Neural Information Processing Systems 26, pages 1250–1258. Curran Associates, Inc.

Pnevmatikakis, E. A., Soudry, D., Gao, Y., Machado, T. A., Merel, J., Pfau, D., Reardon, T., Mu, Y., Lacefield, C., Yang, W., Ahrens, M., Bruno, R., Jessell, T. M., Peterka, D. S., Yuste, R., and Paninski, L. (2016). Simultaneous denoising, deconvolution, and demixing of calcium imaging data. Neuron, 89(2):285–299.

Reynolds, S., Abrahamsson, T., Schuck, R., Sjöström, P. J., Schultz, S. R., and Dragotti, P. L. (2017). ABLE: An Activity-Based level set segmentation algorithm for Two-Photon calcium imaging data. eNeuro, 4(5).

Rupprecht, P., Carta, S., Hoffmann, A., Echizen, M., Blot, A., Kwan, A. C., Dan, Y., Hofer, S. B., Kitamura, K., Helmchen, F., and Friedrich, R. W. (2021). A database and deep learning toolbox for noise-optimized, generalized spike inference from calcium imaging. Nat. Neurosci., 24(9):1324–1337.

Soltanian-Zadeh, S., Sahingur, K., Blau, S., Gong, Y., and Farsiu, S. (2019). Fast and robust active neuron segmentation in two-photon calcium imaging using spatiotemporal deep learning. Proceedings of the National Academy of Sciences, 116(17):8554–8563.

Song, A., Charles, A. S., Koay, S. A., Gauthier, J. L., Thiberge, S. Y., Pillow, J. W., and Tank, D. W. (2017). Volumetric two-photon imaging of neurons using stereoscopy (vtwins). Nature Methods, 14(4):420–426.

Spaen, Q., Asín-Achá, R., Chettih, S. N., Minderer, M., Harvey, C., and Hochbaum, D. S. (2019). Hnccorr: A novel combinatorial approach for cell identification in calcium-imaging movies. eNeuro, 6(2).

Wu, Y., Kirillov, A., Massa, F., Lo, W.-Y., and Girshick, R. (2019). Detectron2. https://github.com/facebookresearch/detectron2.

Xie, M. E., Adam, Y., Fan, L. Z., Böhm, U. L., Kinsella, I., Zhou, D., Paninski, L., and Cohen, A. E. (2020). High fidelity estimates of spikes and subthreshold waveforms from 1-photon voltage imaging in vivo. bioRxiv, 2020.01.26.920256.

Yang, W., Miller, J.-E. K., Carrillo-Reid, L., Pnevmatikakis, E., Paninski, L., Yuste, R., and Peterka, D. S. (2016). Simultaneous multi-plane imaging of neural circuits. Neuron, 89(2):269–284.

Zhang, Y., Zhang, G., Han, X., Wu, J., Li, Z., Li, X., Xiao, G., Xie, H., Fang, L., and Dai, Q. (2022). Rapid deep widefield neuron finder driven by virtual calcium imaging data. bioRxiv.

Zhou, P., Reimer, J., Zhou, D., Pasarkar, A., Kinsella, I., Froudarakis, E., Yatsenko, D. V., Fahey, P. G., Bodor, A., Buchanan, J., Bumbarger, D., Mahalingam, G., Torres, R., Dorkenwald, S., Ih, D., Lee, K., Lu, R., Macrina, T., Wu, J., da Costa, N., Reid, R. C., Tolias, A. S., and Paninski, L. (2020). EASE: EM-Assisted source extraction from calcium imaging data. bioRxiv, 2020.03.25.007468.

Zhou, P., Resendez, S. L., Rodriguez-Romaguera, J., Jimenez, J. C., Neufeld, S. Q., Giovannucci, A., Friedrich, J., Pnevmatikakis, E. A., Stuber, G. D., Hen, R., Kheirbek, M. A., Sabatini, B. L., Kass, R. E., and Paninski, (2018). Efficient and accurate extraction of in vivo calcium signals from microendoscopic video data. eLife, 7:1–14.

